# Biphasic response of CD8 T cell to asparagine restriction maximizes its metabolic fitness and antitumoral functionality

**DOI:** 10.1101/2022.07.18.500458

**Authors:** JN Rashida Gnanaprakasam, Lingling Liu, Xuyong Chen, Siwen Kang, Tingting Wang, Teresa A. Cassel, Christopher M. Adams, Richard M Higashi, David A. Scott, Gang Xin, Jun Yang, Andrew N. Lane, Teresa W.-M. Fan, Ji Zhang, Ruoning Wang

**Author notes:** Correspondence should be addressed to: Ruoning Wang.

## Abstract

Robust and effective T cell immune surveillance and cancer immunotherapy require properly allocating metabolic resources to sustain energetically costly processes, including growth and cytokine production. Amino acids are major cellular constituents that serve as protein building blocks, energy sources, and signaling molecules. Although T cells can synthesize all nonessential amino acids, including asparagine (Asn), activated CD8 T cells still consume considerable quantities of exogenous Asn. Unexpectedly, Asn restriction on CD8 T cells induced a biphasic response, consisting of sequential actions with opposing effects at two conceptually separated phases after activation. Asn restriction suppressed activation and cell cycle entry in the early phase by depleting the intracellular Asn pool while rapidly engaging an ATF4/NRF2-dependent stress response, conferring robust proliferation and effector function of CD8 T cells in the late phase. Mechanistically, ATF4 and NRF2 activation rendered CD8 T cells to utilize de novo biosynthesis of Asn, consuming less glucose and glutamine but producing more intracellular nucleotides for proliferation. Moreover, NRF2 activation promoted the expression of inflammatory and effector genes to enhance effector functions in CD8 T cells. Accordingly, Asn restriction or overexpression of ATF4 or NRF2 potentiated T cell-mediated antitumoral response in the metabolically restricted tumor microenvironment. Our studies revealed Asn as a critical metabolic node in directing the stress signaling to shape T cell metabolic fitness and effector functions. Asn restriction is a promising and clinically relevant strategy to enhance cancer immunotherapy.

## Introduction

T cell response’s exquisite specificity, amplitude, and quality are critical for immune surveillance and cancer immunotherapy. However, T cell response is a metabolically costly process. It is often constrained by the metabolic landscape of the tissue microenvironment and the rapidly proliferating pathogens that compete for nutrient resources with the host. T cell activation rapidly engages the central carbon catabolic pathways, including glycolysis, the pentose phosphate pathway (PPP), and the tricarboxylic acid (TCA) cycle, to prepare cells for growth, differentiation, and immune defense (*1–4*). An optimal allocation of limited energy and nutrient resources between growth, repair, and production of effector molecules is required to maximize T cell-mediated responses. The largest constituent of cell mass comes from amino acids in proteins (*5*). Amino acids are building blocks of protein, carbon/nitrogen sources for energy, anabolic substrates, and signaling molecules (*6*). T cells cannot synthesize essential amino acids like other mammalian cells, thus relying on exogenous sources. Although T cells can synthesize all the nonessential amino acids (NEAA), a robust proliferation and effector response requires the exogenous supply of several NEAAs (*7–9*). The bioavailability of amino acids is tightly coupled with the central carbon metabolism and cell signaling transduction that shape T cell proliferation, differentiation, and immunological functions (*10, 11*). CD8 effector T (T_eff_) cells play a major role in antitumor immunity and elicit antitumor activity through direct recognition and killing of antigen-presenting tumor cells and orchestrating numerous adaptive and innate immune responses (*12–15*). However, tumors can co-opt various immunosuppressive mechanisms that act in concert to foster an immune-tolerant microenvironment and thus escape the T cell-mediated antitumor immune response (*16–19*). As activated T cells share metabolic characteristics with tumor cells (*1, 20*) the metabolically demanding cancer cells restrict the function of effector T cells by competing for nutrients and producing immunosuppressive metabolites (*16, 21*). The development of CD8 T_eff_ responses can be broadly classified into two distinct and sequential phases—an initial activation phase in which CD8 T cells accumulate cellular mass and prepare to divide, followed by the differentiation phase whereby CD8 T cells rapidly expand and differentiate into effector cells (*22*). Shifts in energy and carbon expenditure accompany the early to late phase transition to support different biologic activities and effector functions. A better understanding of the metabolic dependence of CD8 T cells and the metabolic interplay within the tumor microenvironment (TME) will enable us to devise rational and effective approaches to improve T cell metabolic fitness and cancer immunotherapy.

## Results

### Metabolic profiling reveals that CD8 effector T cells differentially consume and depend on extracellular nonessential amino acids

To assess how the bioavailability of amino acids and their utilization impact CD8 T cells at two conceptually separated phases, we measured the overall consumption and excretion rate of each amino acid in cells in the early activation and late effector phases by using GC/MS analysis (**Fig. 1A**). We also determined the extent of intracellular amino acid accumulation in cells (**Fig. S1A**). CD8 cells at both phases displayed higher levels of intracellular amino acids than naïve CD8 T cells (**Fig. S1A**). Like other mammalian cells, T cells obtained all the essential amino acids (EAA) from the extracellular environment (cell media). Although T cells can synthesize all the nonessential amino acids (NEAA), CD8 T cells still consumed a significant amount of glutamine (Gln), asparagine (Asn), arginine (Arg), and serine (Ser) after activation (**Fig. 1A**). Proline (Pro) and tyrosine (Tyr) were consumed in the early phase and the late phase, respectively (**Fig. 1A**). The rest of the NEAAs were excreted into the media by CD8 T cells in both phases after activation (**Fig. 1A**).

**Figure 1.**
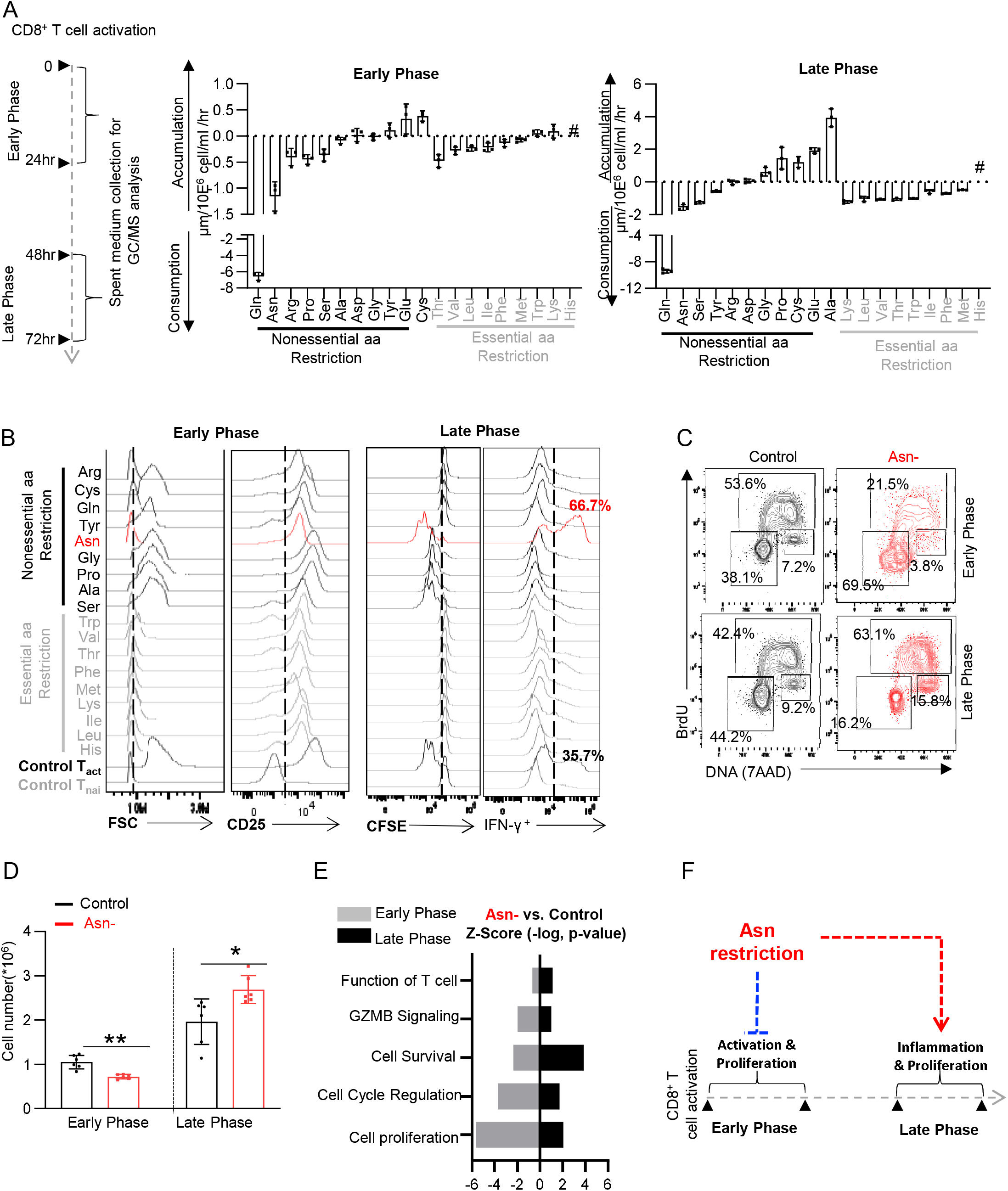
Asn restriction on CD8 T cells induces a biphasic response. A) Schematic diagram of measuring amino acid consumption and production of T cells in the early (24hr) and late phase (72hr) after activation (left). Indicated amino acids in the culture medium collected from T cells in the early (middle), and the late phase (right) were quantified by GC-MS. The consumption (negative value) or production (positive value) rate of indicated amino acids was determined by calculating the difference between spent and blank medium. # Indicates amino acids that were not quantified due to technical limitations. Arginine was quantified by an arginine assay kit (n=3). Data are representative of 3 independent experiments. B) CD8 T cells were activated in a complete medium or indicated single amino acid-deficient medium. Cell size (FSC) and cell surface expression of CD25 were determined by flow cytometry in the early phase (24hr). Proliferation (CFSE) and intercellular expression of IFN-γ were determined by flow cytometry in the late phase (72hr). Flow plots are representative of 2 independent experiments. C-E) CD8 T cells were activated in a medium with or without Asn and were collected in the early phase (24-36hr) and late phase (72hr). The cell cycle profile (C) was determined by flow cytometry. The numbers indicate the percentage of cells in the cell cycle stage (n=3); Data are representative of 3 independent experiments. The cell number (D) was determined by a cell counter (n=5). Data are representative of 3 independent experiments. Error bars are mean ±SD. *p < 0.05, **p < 0.01, Student’s t-test. Comparative differently expressed gene signatures (E) were determined by the IPA analysis of RNAseq (24hr) and listed according to their z score. F) The conceptual model of a biphasic response induced by Asn restriction.

We reasoned that CD8 T cells might differentially rely on extracellular NEAAs for in vitro activation, proliferation, and effector function as some were consumed from the media, and others were excreted by cells. To systematically assess the amino acid dependence of T cells, we removed each of the 18 amino acids from culture media and comprehensively profiled cell activation markers, size, proliferation, and IFN-γ production. Of note, glutamate (Glu) or aspartate (Asp) restriction were excluded from the experiment because non-neuronal cell types, including T cells, cannot transport these two amino acids (*23*). The activation marker CD25 was induced after activation under any single amino acid restriction condition, albeit to various degrees (**Fig. 1B**). However, cell growth in the early phase, proliferation, and cytokine production in the late phase strictly require all essential and several nonessential amino acids, including Gln, Arg, Cys, and Tyr (**Fig. 1B**). These findings align with the known role of NEAAs in supporting T cell bioenergetic and biosynthetic activities (*11, 24–33*). Intriguingly, Asn restriction remarkably reduced cell size and the level of activation marker in the early phase but enhanced proliferation and the percentage of IFN-γ^+^ CD8 T_eff_ cell in the late phase (**Fig. 1B**), indicating that Asn restriction elicited opposing effects on CD8 T cells in the early and the late phase after activation.

### Asn restriction on CD8 T cells induces a biphasic response

Next, we comprehensively assessed the effects of Asn restriction on CD8 T cells in the early and late phases after activation. In the early phase, Asn restriction significantly delayed cell cycle progression from G0/1 to S phase (**Fig.1C**), reduced cell numbers and protein content (**Fig.1D** **and S1D**), which were associated with moderately reduced cell RNA/DNA content and cell viability (**Fig. S1B-C**). In contrast, Asn restriction increased the percentage of cells in the S-phase and increased total cell numbers without affecting the level of protein/DNA/RNA content and cell viability in the late phase (**Fig. 1C-D** **and S1B-D**). Asparaginase (L-ASP) has been used in the clinic to treat acute lymphoblastic leukemia (ALL) by depleting Asn in the blood (*34, 35*). We confirmed that recombinant L-ASP treatment increased cell numbers and the percentage of IFN-γ^+^ CD8 T_eff_ cells in the late phase, mirroring the effects of Asn-free conditional media on CD8 T cells (**Fig. S1E-F**). We then assessed the homeostatic proliferation of CD8 T cells in vivo following the treatment of L-ASP with a sufficient dose to deplete serum Asn and increase serum aspartate in mice (**Fig. S1G-H**). In agreement with our in vitro data, L-ASP-mediated depletion of Asn in serum did not affect adoptively transferred CD8 T cell homeostatic proliferation (**Fig. S1I**).

To gain mechanistic insights into the effects of Asn restriction on CD8 T cells, we performed RNA-seq in cells collected in the early and late phases, which revealed opposing gene enrichment patterns for the genes involved in cell cycle/proliferation, cell survival, and T cell function/signaling in the early phase versus the late phase (**Fig. 1E, S2A-C and S3A-C**). In the early phase, Asn restriction suppressed the expression of T cell activation genes and most of the genes that promote cell cycle progression but enhanced the expression of genes involved in cell cycle checkpoint response and thus suppressed cell cycle progression (**Fig. S2A-C**). By contrast, CD8 T_eff_ cells in the late phase expressed higher levels of cell cycle-promoting genes, inflammatory signaling, and cytokine genes but lower levels of cell cycle checkpoint genes under Asn restriction than in the control condition (**Fig. S3A-C**). Our data suggested that Asn restriction induced a biphasic response and conferred robust proliferation and effector function to CD8 T cells in the late phase (**Fig. 1F**).

### Asn restriction enhances the CD8 T cell-mediated antitumor responses in vitro and in vivo

The finding that Asn restriction significantly enhanced T_eff_ cell proliferation and inflammatory cytokine production prompted a closer examination of the role of Asn in CD8 T_eff_ cell-mediated antitumoral response. We generated antigen-specific mouse T_eff_ cells using a major histocompatibility complex (MHC) class I-restricted premelanosome protein (Pmel)-1 T cell receptor model (**Fig. 2A**) (*36–38*). We also generated disialoganglioside (GD2)-specific chimeric antigen receptor (GD2-CAR) human T_eff_ cells (**Fig. 2A**) (*39*). Pmel T_eff_ cells recognized Pmel-17 (mouse homologue of human SIVL/gp100) in mouse melanoma, while GD2-CAR T cells recognized GD2 expressed in tumors with neuroectodermal origin. Asn restriction increased the expression and production of proinflammatory cytokines (tumor necrosis factor/TNFα and IFN-γ) and effector molecule (granzyme b/GZMB) in Pmel T_eff_ cells and GD2-CAR T cells (**Fig. 2B** **and S4A-D**). In addition, Asn restriction increased the cell number in these two models (**Fig. 2C**). We then co-cultured Pmel T_eff_ cells with Pmel^+^ B16F10-gp100 mouse melanoma cells and GD2-CAR T cells with GD2^+^ LAN-1 human neuroblastoma cells, respectively. Asn restriction significantly enhanced the tumor-killing activities of T cells in both models (**Fig. 2D**). Notably, B16F10 and Lan-1 are Asn-prototrophic cell lines that could grow in Asn-free media in vitro (**Fig. S4E-H**).

**Figure 2.**
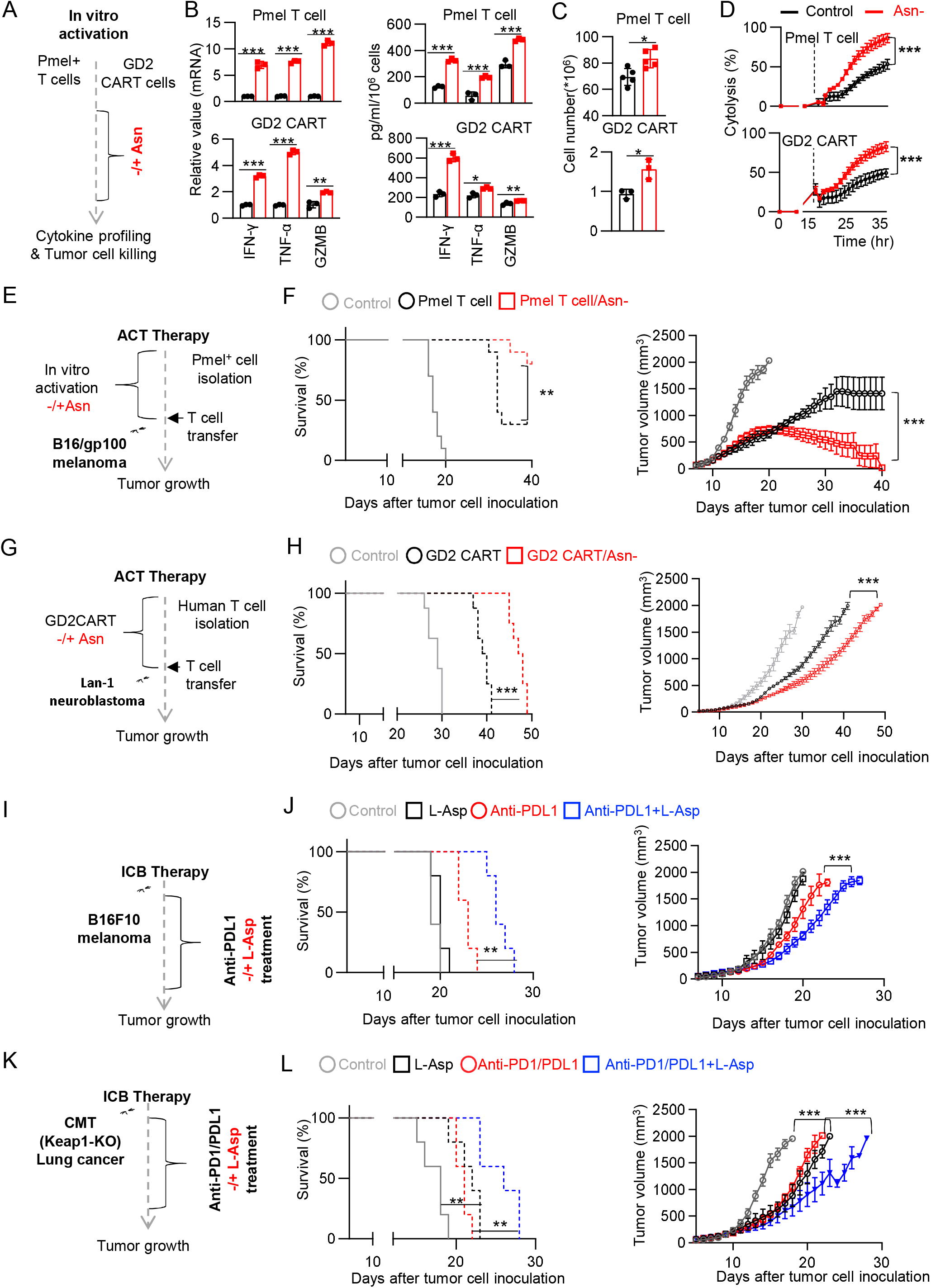
Asn restriction enhances the T cell-mediated antitumor responses. A-D) Schematic diagram of in vitro assays (A). The level of indicated mRNAs in Pmel T cells (upper) and GD2 CART (lower) were determined by qPCR. Indicated proteins in culture supernatants were quantified by ELISA and legend plex (B) (n=3). The cell number was determined by a cell counter (C) (n=3-5). The cytolysis was determined by eSight (D); (n=3). Data from A-D are representative of 3 independent experiments. Error bars are mean ±SD. *p < 0.05, **p < 0.01, *** P < 0.001, Student’s t-test. E-L) Schematic diagrams of animal experiments (E, G, I, and K). Kaplan-Meier survival curves and tumor growth of mice bearing B16-gp100 tumor (n=10) (F). Data are representative of 3 independent experiments. Kaplan-Meier survival curves and tumor growth of mice bearing LAN-1 neuroblastoma tumor (n=8) (H). Data are representative of 2 independent experiments. Kaplan-Meier survival curves and tumor growth of mice bearing B16F10 melanoma tumor (n=5) (J). Data are representative of 3 independent experiments. Kaplan-Meier survival curves and tumor growth of mice bearing CMT Keap1 KO lung tumor (n=5) (L). Data are representative of 2 independent experiments. Error bars represent mean ± SEM (F, H, J, and L). **p < 0.01, *** P < 0.001, Two-way ANOVA for tumor growth curve and log-rank test for animal survival.

The adoptive cell transfer (ACT) of tumor-infiltrating lymphocytes or CAR T cells and the checkpoint blockade (ICB) based on monoclonal antibodies targeting immune checkpoint proteins are front-line cancer immunotherapies. We reasoned that Asn restriction might improve the efficacies of cancer immunotherapies that could engage CD8 T cell antitumoral activities. We expanded Pmel T cells and GD2-CAR T cells with or without Asn in vitro, followed by transferring Pmel T cells to mice bearing B16 melanoma xenografts (**Fig. 2E-F** **and S4L**) and GD2-CAR T cells to immune-deficient mice bearing GD2 positive human neuroblastoma (LAN-1) xenografts (**Fig. 2G-H** **and S4M**). While adoptively transferred T cells alone significantly delayed tumor growth and prolonged animal survival time, expanding Pmel+ or GD2-CAR T cells under Asn restriction further potentiated their tumor restraining effects in these two ACT models (**Fig. 2E-H**). PD1 and PDL1 blockade has been shown to elicit durable T cell-dependent antitumor responses in melanoma patients and a pre-clinical model of melanoma (B16 melanoma-bearing mice) (*40, 41*). We thus assessed the antitumor effect of combining L-ASP and anti-PDL1 antibodies in B16 melanoma-bearing mice. In agreement with our in vitro data (**Fig. S4E**), L-ASP-mediated depletion of Asn did not affect tumor cell growth in vivo (**Fig. 2I-J** **and S4N**). Moreover, mice treated with L-ASP and anti-PDL1 antibodies displayed a better outcome than the anti-PDL1 monotherapy group (**Fig. 2I-J**). We also examined the effect of combining Asn restriction by L-Asp treatment and ICB (anti-PD1/L1 antibodies) in a pre-clinical model of lung carcinoma (CMT167-bearing mice) (*42*). Kelch-like ECH-associated protein 1 gene (Keap-1) is one of the most frequently mutated tumor suppressor genes in non-small cell lung cancer (NSCLC), and the loss of its function renders cancer cells sensitive to NEAAs (including Asn) restriction (*43, 44*). We reasoned that a Keap-1 deficient tumor model could allow us to test L-ASP monotherapy and ICB combination therapy. Thus, we generated a Keap-1 KO CMT cell line by CRISPR and confirmed that Asn restriction decreased cell proliferation and viability in vitro (**Fig. S4I-K**). While either L-Asp or ICB alone reduced tumor growth and prolonged animal survival time, the combinational therapy further potentiated their antitumor effects (**Fig. 2K-L** **and S4O**). These results indicated that Asn restriction might be a clinically relevant strategy for enhancing cancer immunotherapies.

### Asn restriction promotes Asn de novo biosynthesis in the late but not the early phase

Next, we sought to ask why exogenous Asn is dispensable for driving T cell growth in the early phase but becomes indispensable for cell proliferation in the late phase. We envisioned that the CD8 T_eff_ cell might engage de novo biosynthesis of Asn through Asn synthetase (ASNS) to support T cell proliferation in the late phase (**Fig. S5A**) (*45, 46*). Supporting this hypothesis, activation of T cells under Asn restriction led to a time-dependent upregulation of ASNS, in line with a partial restoration of intracellular Asn pool in the late but not the early phase after T cell activation (**Fig. S5B-C**). As Gln is a vital carbon donor for replenishing the TCA cycle metabolites and Asp biosynthesis, the fraction of ^13^C_5_ and ^13^C_4_ isotopologues (generated via oxidation of ^13^C_5_-Gln) of the TCA cycle intermediate metabolites and Asn were significantly increased in the context of Asn restriction in the late phase (**Fig. S5D**).

Next, we generated a T cell-specific ASNS knockout mouse strain (ASNS KO) by crossing the ASNS^fl^ mouse strain with the CD4-Cre mouse strain. ASNS deletion did not result in any notable defects in T cell development in the thymus, spleen, and lymph node (**Fig. S6A**). Immune blot (IB) analyses validated the deletion of ASNS in T cells (**Fig. 3A**), which depleted the intracellular Asn pool under Asn restriction (**Fig. 3B-C**). In the early phase after activation, ASNS deletion did not change the effects of Asn restriction on CD8 T cell activation or size (**Fig. 3D**). While ASNS deletion alone did not affect CD8 T_eff_ cell proliferation, it abolished CD8 T_eff_ cell proliferation and inflammatory cytokine production under Asn restriction (**Fig. 3E-G**). L-ASP-mediated depletion of Asn significantly dampened homeostatic proliferation in adoptively transferred ASNS-KO but not WT CD8 T cells, confirming that ASNS activity rendered T cells independent of exogenous Asn in vitro and in mice (**Fig. 3H-J**).

**Figure 3.**
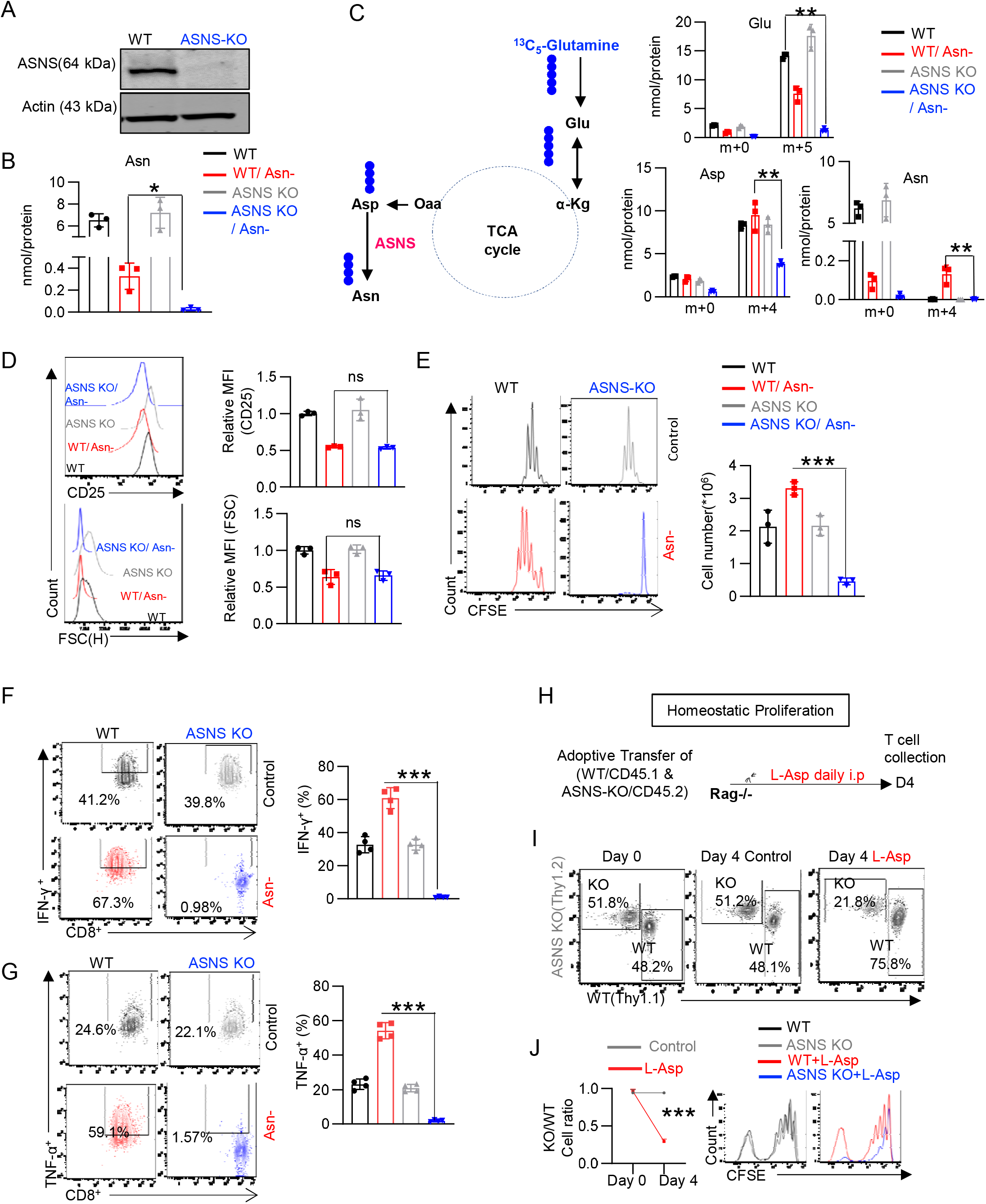
Asn restriction renders T cell dependent on ASNS expression. A) The indicated protein was determined by immunoblot. B) The level of Asn in the indicated groups was determined by GC-MS (n=3). C) Diagram of [^13^C_5_]-Glutamine catabolism through entering the downstream TCA cycle and Asn biosynthesis. ●: denoted the ^13^C label of all carbons of indicated metabolites derived from [^13^C_5_]-glutamine catabolism (left panel). Metabolites in the indicated groups were analyzed by GC-MS (right panel), numbers on the X-axis represent those of ^13^C atoms in given metabolites, and numbers on the Y-axis represent the levels of the metabolites (nmol/ protein). Glu: glutamate, Asp: aspartate, Asn: asparagine, (n=3). Data are representative of one experiment. D-G) CD8 T cells with the indicated genotype were activated in a medium with or without Asn and collected at 24hr (the early phase, D) and 72hr (the late phase, E-G). Cell surface expression of CD25 (top, D), cell size (FSC) (bottom, D) (n=3), proliferation (CFSE) (E) (n=3), and intercellular expression of indicated proteins (F-G) (n=4) were determined by flow cytometry. Cell number (E) was determined by a cell counter (n=3). Data from D-G are representative of 3 independent experiments. H-J) Schematic diagram of in vivo competitive proliferation (H). The donor cell ratios before and after adoptive transfer were determined by surface staining of isogenic markers (I). Cell proliferation was determined by CFSE dilution (J) (n=3). Data are representative of 3 independent experiments. Error bars represent mean ± SD. *p < 0.05, **p < 0.01, *** P < 0.001, ns, not significant, Student’s t-test.

### Asn restriction-induced an ATF4-mediated response in CD8 T cells

Next, we investigated the mechanism of how ASNS is activated by Asn restriction. RNAseq analysis of CD8 T cells in the early phase revealed several enriched gene signatures under Asn restriction that are regulated by activating transcription factor 4 (ATF4) and nuclear factor-erythroid factor 2-related factor 2 (NRF2) (**Fig. 4A**). ASNS was revealed as a direct transcriptional target of ATF4 in mammalian cells, which mediates an integrated stress response upon a range of environmental insults, including nutrient restriction and oxidative stress (*47*). Asn restriction in CD8 T cells increased the phosphorylation of general control nonrepressed 2 (GCN2), the upstream regulator of ATF4, and the expression of ATF4 and NRF2 as early as 4hr after activation (**Fig. 4B-C**). Next, we generated a T cell-specific ATF4 knockout mouse strain (ATF4 KO) by crossing the ATF4^fl^ mouse strain with the CD4-Cre mouse strain. ATF4 deletion did not result in defects in T cell development in the thymus, spleen, and lymph node (**Fig. S7A**). In line with the previous report that ATF4 controlled the expression of ASNS and mediated oxidative stress response (*47*), deleting ATF4 abolished the expression of ASNS and NRF2 in CD8 T cells under Asn restriction (**Fig. 4D-E**), raising the possibility that both ATF4 and NRF2 are involved in regulating T cell proliferation and inflammation upon Asn restriction (**Fig. 4F**). In the early phase after activation, ATF4 deletion did not change the effects of Asn restriction on CD8 T cell activation and size (**Fig. S7B**). However, in the late phase, ATF4 KO CD8 T_eff_ cell failed to proliferate, maintain survival and produce inflammatory cytokine under Asn restriction (**Fig. 4G-I** **and S7C**).

**Figure 4.**
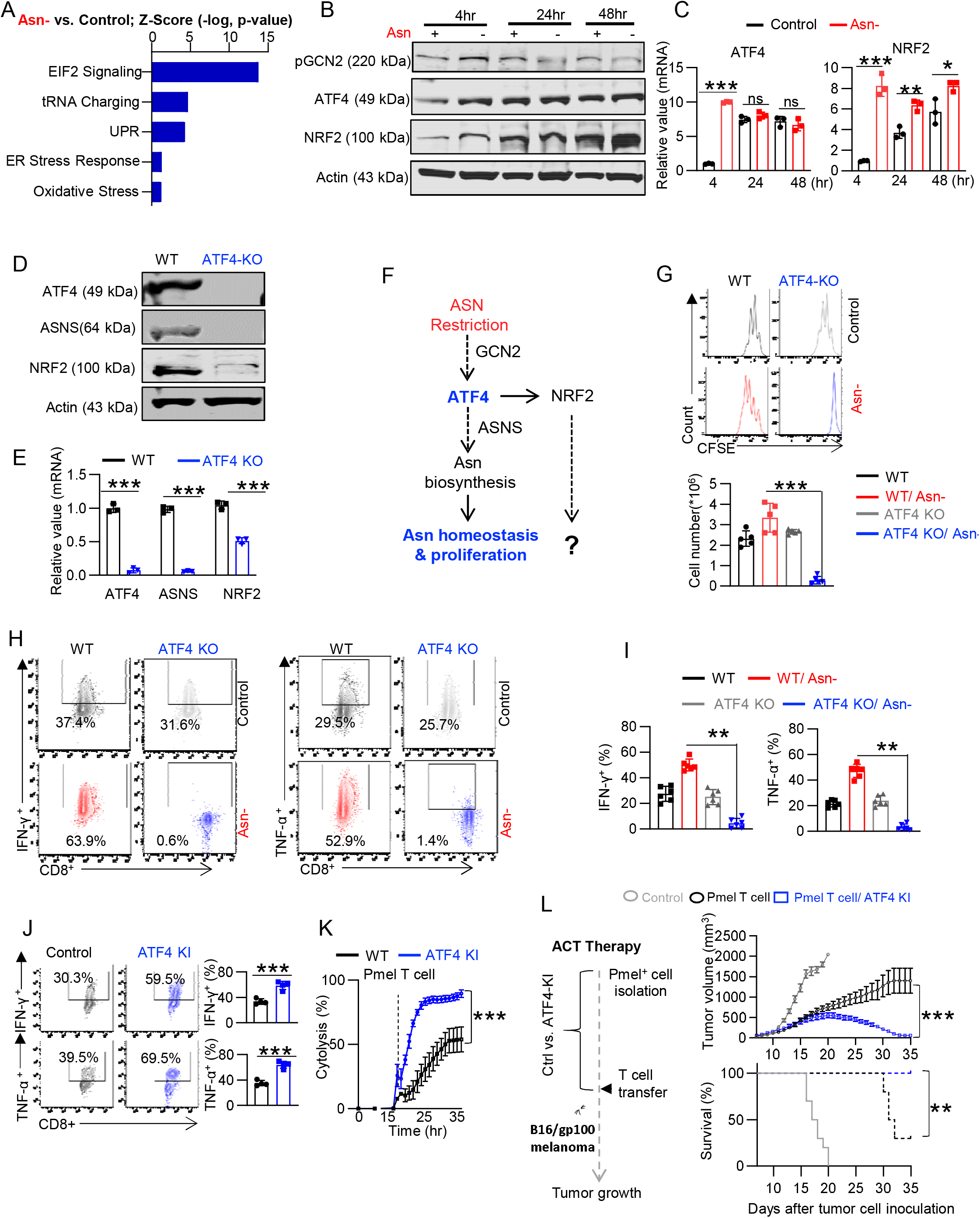
Asn restriction induces ATF4 expression in the early phase that enhances the anti-tumor response in the late phase. A) Differently expressed gene signatures were determined by the IPA analysis of RNAseq (cells collected 4hr after activation) and listed according to their z score. Data is representative of one experiment. B-C) CD8 T cells were activated in a medium with or without Asn for the indicated time. Protein and mRNA levels of indicated molecules were determined by Immunoblot (B) and qPCR (C) (n=3). Data are representative of 2 independent experiments. D) CD8 T cells with the indicated genotype were collected 4hr after activation in a medium without Asn. The level of indicated protein and mRNA was determined by immunoblot (D) and qPCR (E) (n=3). Data are representative of 2 independent experiments. F) Schematic conceptual model of ATF4 and possibility of NRF2 in regulating T cell proliferation under Asn restriction. G-I) CD8 T cells from indicated genotype were activated in a medium with or without Asn for 72hr. Cell proliferation was determined by CFSE staining, and the cell number was determined by a cell counter (G) (n=5). Data are representative of 4 independent experiments. The expression of indicated cytokines was determined by intracellular staining by flow cytometry (H-I) (n=6). Data are representative of 3 independent experiments. Error bars represent mean ± SD. J-L) Cells with indicated genotypes were activated for 5 days, followed by restimulation with soluble antibodies (2 days). The expression of indicated cytokines was determined by flow cytometry (J) (n=3). Data are representative of 3 independent experiments. The cytolysis was determined by eSight (K) (n=3). Data are representative of 2 independent experiments. Error bars represent mean ± SD. Schematic diagram of the experiment (left, L). Tumor growth (top) and Kaplan-Meier survival curves (bottom) of mice bearing B16-gp100 tumor (right, L) (n=10). Data are representative of 2 independent experiments. Error bars represent mean ± SEM. *p < 0.05, **p < 0.01, *** P < 0.001, ns, not significant, Student’s t-test, Two-way ANOVA for tumor growth curve, and log-rank test for animal survival.

Next, we sought to determine if ATF4 overexpression could resemble the effects of Asn restriction on CD8 T_eff_ cell proliferation and function. We crossed the ATF4^LSL-KI^ strain with the CD4-Cre strain and the Pmel transgenic strain to generate a Pmel^+^ ATF4-KI strain to assess the effect of ATF4 overexpression in T cells in an antigen-specific manner. Compared to the WT control cells, Pmel^+^ ATF4-KI T_eff_ cells expressed a higher level of inflammatory cytokines and were more effective in killing tumor cells in vitro and controlling tumor growth in mice (**Fig. 4J-L** **and S7D-F**). These results indicated that ATF4-dependent ASNS expression enabled T cell proliferation under Asn restriction. Notably, ATF4 overexpression could phenocopy the effects of Asn restriction on T cells by inducing inflammation.

### NRF2 is required for enhancing proliferation and inflammation by Asn restriction

We have shown that ATF4-dependent ASNS expression is required for proliferation and cytokine production under Asn restriction (**Fig. 4D and G**). However, we reasoned that cytokine production defects might be due to the lack of proliferation in ATF KO CD8 T cells. Since ATF4 is required for inducing NRF2 expression upon Asn restriction (**Fig. 4D**), we sought to determine the role of NRF2 in regulating CD8 T_eff_ cells by using a germline *Nrf2* knockout mouse strain (*48*). NRF2 deletion did not affect the expression of ASNS and ATF4 in CD8 T cells (**Fig. 5A**) and therefore did not abolish proliferation under Asn restriction (**Fig. 5B**). However, NRF2 deletion alleviated the hyperproliferation phenotype of CD8 T cells induced by Asn restriction (**Fig. 5B**). Importantly, Asn restriction failed to enhance the expression of inflammatory cytokines and transcription factors in NRF KO but not WT CD8 T cells (**Fig. 5C-E**). Next, we assessed the effect of NRF2 gain-of-function on Pmel^+^ CD8 T_eff_ cells by retroviral transduction of a dominant-active form of NRF2 (NRF2-OE) (*49*). NRF2-OE phenocopied the effects of Asn restriction on T cells by increasing the expression of inflammatory cytokines, killing tumor cells more effectively in vitro, and potentiating the antitumoral effects in mice (**Fig. 5F-H** **and S8A-C**). Collectively, our results suggested that the action of ATF4 activation on CD8 T cells is bimodal, consisting of the sequential action of two discrete processes, ASNS-dependent Asn biosynthesis and NRF2-mediated effects on enhancing proliferation and effector functions (**Fig. 5I**).

**Figure 5.**
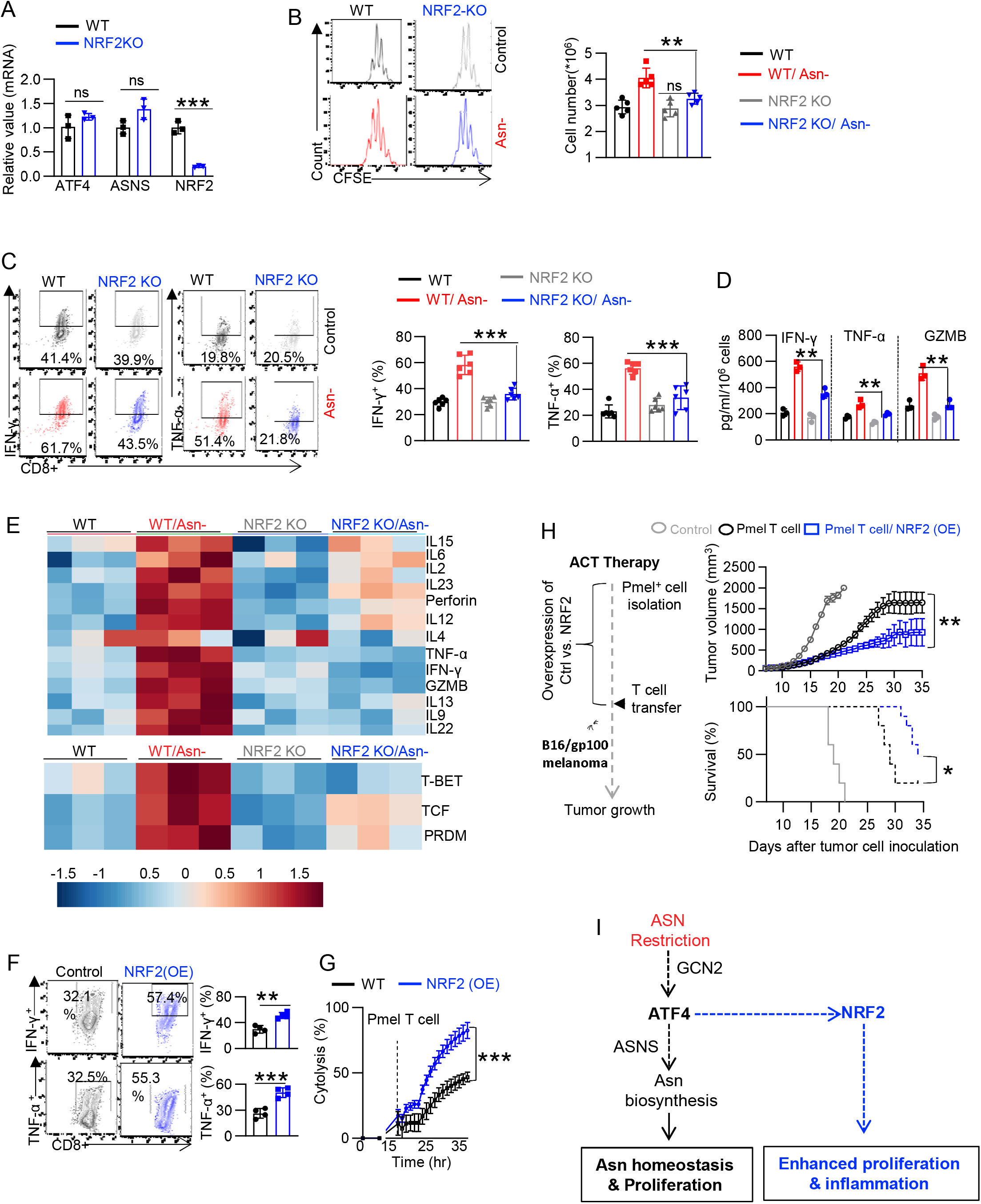
NRF2 is required for enhancing proliferation and inflammation by Asn restriction. A) The level of the indicated mRNA in cells collected 4hr after activation was determined by qPCR (n=3). Data are representative of 2 independent experiments. B-E) CD8 T cells from indicated genotypes were activated in a medium with or without Asn for 72hr. Cell proliferation was determined by CFSE staining, and cell number was determined by a cell counter (B) (n=5). Data are representative of 4 independent experiments. The expression of indicated cytokines was determined by intracellular staining by flow cytometry (C) (n=6). Data are representative of 3 independent experiments. The level of indicated proteins in the culture medium collected 72hr after activation was quantified by ELISA (D) (n=3). Data are representative of 2 independent experiments. The level of the indicated mRNA in cells collected 72hr after activation was determined by qPCR and depicted by heatmap (E) (n=3). F-H) Cells with indicated genotypes were activated for 5 days, followed by restimulation with soluble antibodies (2 days). The expression of indicated cytokines was determined by flow cytometry (F) (n=3). Data are representative of 3 independent experiments. The cytolysis was determined by eSight (G) (n=3). Data are representative of 2 independent experiments. Error bars represent mean ± SD. Schematic diagram of the experiment (left, H). Tumor growth (top) and Kaplan-Meier survival curves (bottom) of mice bearing B16-gp100 tumor (right, H) (n=10). Data are representative of 2 independent experiments. Error bars represent mean ± SEM. **p < 0.01, *** P < 0.001, ns, not significant, Student’s t-test, Two-way ANOVA for tumor growth curve, and log-rank test for animal survival.

### Asn restriction optimizes carbon assimilation to increase nucleotide bioavailability in the late phase

We have shown that CD8 T cells could engage ASNS-dependent de novo synthesis to partially replenish the intracellular Asn pool under Asn restriction (**Fig. S5A and 3C**). However, the level of intracellular Asn pool under Asn restriction was still lower than the control group in the late phase, indicating that engaging ASNS-dependent de novo synthesis could not completely restore the Asn pool (**Fig. S5C**). We, therefore, reasoned that in addition to engaging Asn de novo biosynthesis, Asn restriction might cause more changes in the central carbon metabolism to sustain a robust proliferation in the late phase. To test this hypothesis, we took an integrated stepwise approach to comprehensively analyze the metabolic profiles of CD8 T cells in the late phase (**Fig. 6A**). The extracellular metabolome reflects the ultimate outcome of metabolic input, processing, and output. We first assessed the overall consumption rate of glucose and glutamine, two primary carbon sources for proliferating cells. Under Asn restriction, CD8 T_eff_ cells consumed much less glucose and glutamine and excreted less lactate and glutamate than the control group (**Fig. 6B**). While Asn restriction reduced the metabolic rate of glycolysis, it increased the metabolic rate of the pentose phosphate pathway (PPP) (**Fig. 6C**). Moreover, CD8 T_eff_ cells under Asn restriction displayed a much higher basal oxygen consumption rate (OCR), and increased intracellular ATP production, accompanied by increased mitochondrial mass and mitochondrial DNA content (**Fig. 6D-F**).

**Figure 6.**
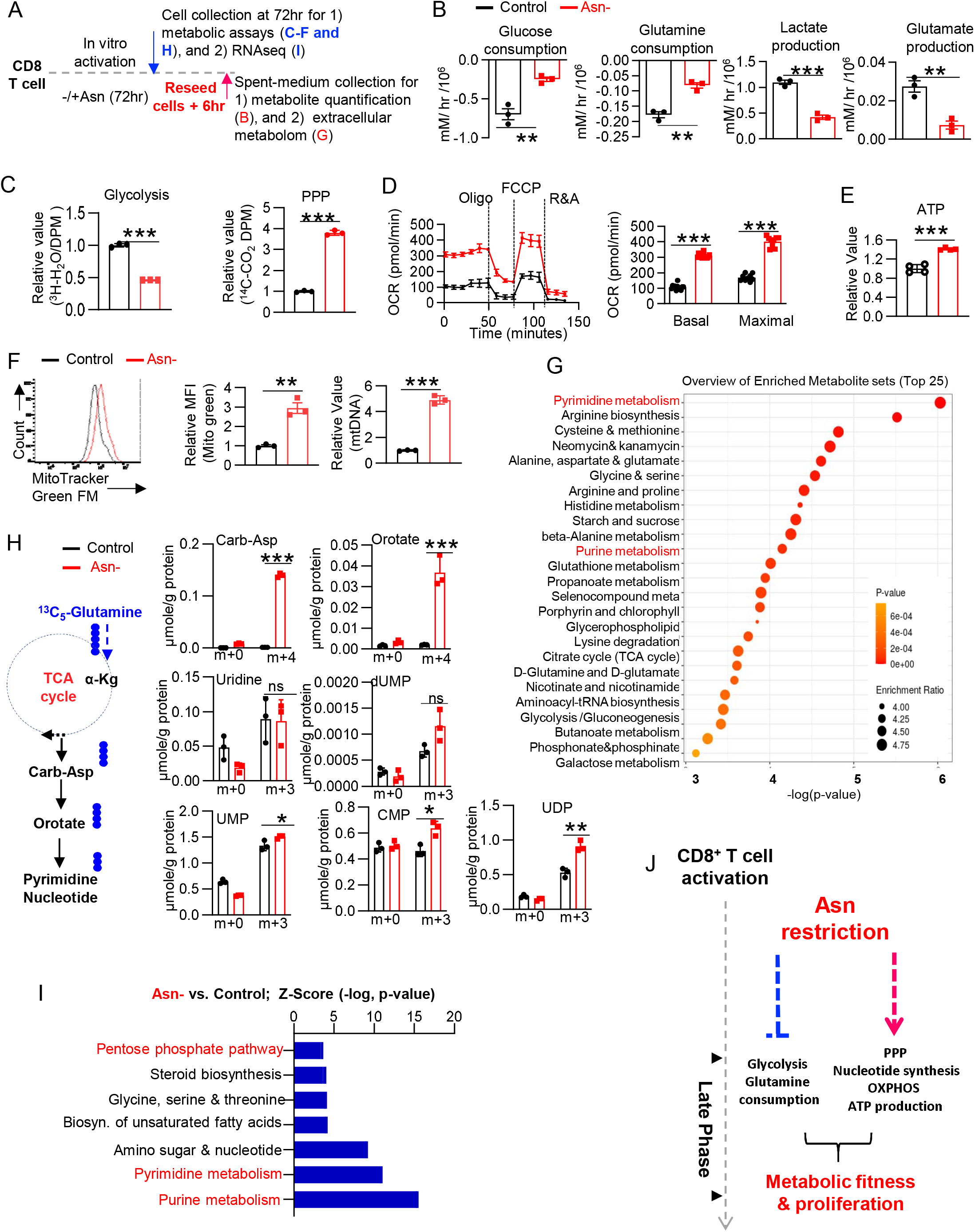
Asn restriction optimizes carbon assimilation to support nucleotides biosynthesis. A) Schematic diagram of sample collection and assays. B) The indicated metabolites were quantified by the YSI bioanalyzer. The consumption and production were determined by calculating the difference between blank and spent medium (6hr incubation of T cells collected in the late phase/72hr) (n=3). C) The glycolysis activity and PPP activity (in the late phase/72hr) were determined by measuring ^3^H_2_O generated from D-[5-^3^H(N)]-glucose and ^14^CO_2_ generated from [1-^14^C] D-glucose, respectively (n=3). D) OCR (in the late phase/72hr) was determined by Seahorse (n=3). E) ATP levels (in the late phase/72hr) were determined by the CellTiter-Glo^®^ 2.0 Assay kit (n=3). F) Mitochondria mass was determined by Mito Tracker green FM staining by flow cytometry, and mitochondrial DNA was quantified by qPCR (n=3). Data from B-F are representative of 3 independent experiments. G) KEGG enrichment analysis of the changes of extracellular metabolites (6hr spent-medium). H) Diagram of [^13^C_5_]-Glutamine catabolism through entering the downstream TCA cycle and pyrimidine biosynthesis. ●: denoted the ^13^C label of all carbons of indicated metabolites derived from [^13^C_5_]-glutamine catabolism (left panel). Metabolites in cells (in the late phase/72hr) were analyzed by IC-UHR-FTMS (right panel), numbers on the X-axis represent those of ^13^C atoms in given metabolites, and numbers on the Y-axis represent the levels of the metabolites (μmole/g protein). Carb-Asp: N-carbamoyl-L-aspartate; UMP/UDP: uridine mono/diphosphate; dUMP deoxyuridine monophosphate; CMP/CDP: cytidine mono/diphosphate (n=3). Data are representative of 2 independent experiments. I) Differently expressed metabolic gene signatures were determined by the IPA analysis of RNAseq (in the late phase/72h). Data from G-I are representative of one experiment. Error bars represent mean ± SD. *p < 0.05, **p < 0.01, *** P < 0.001, ns., not significant, Student’s t-test.

Next, we employed a semi-quantitive untargeted global metabolomics platform to profile metabolites in control (blank) media and the spent media of T cells. In parallel, we collected cells to perform RNAseq (**Fig. 6A**). The principal component analysis revealed that CD8 T_eff_ cells generated in the presence and absence of Asn are characterized by distinct extracellular metabolome profiles (**Fig. S9A**). We further classified metabolites as having changes in consumption or excretion according to whether the level in the spent media is lower or higher than in the blank media. While Asn restriction did not change the overall consumption pattern, it reduced the levels of metabolites that were excreted into the media (**Fig. S9B**). Enrichment analysis revealed that nucleotides and their precursors were among the most differentially changed metabolite groups (**Fig. 6G** **and S9C**). In addition to acting on reducing nucleotide excretion, we reasoned that Asn restriction might increase de novo pyrimidine nucleotide biosynthesis. To test this idea, we supplied ^13^C_5_-glutamine as a metabolic tracer in T cell culture media and followed ^13^C incorporation into downstream metabolites. The corresponding ^13^C_4_ or ^3^C_3_ isotopologues of pyrimidine nucleotides and their precursors were increased under Asn restriction (**Fig. 6H**). Moreover, these metabolic changes were accompanied by related gene expression changes in the PPP and nucleotide de novo biosynthesis, as revealed by RNAseq (**Fig. 6I** **and S10A-B**). Our results suggested that Asn restriction improved T_eff_ cell metabolic fitness by reducing the overall carbon consumption but increasing the ATP production and intracellular nucleotide pool to enhance CD8 T cell proliferation. Optimizing carbon assimilation by reducing the overall carbon consumption might prepare CD8 T cells to function better in the metabolically restricted tumor microenvironment.

### NRF2 is required for rewiring the central carbon metabolism following Asn restriction

NRF2 is critical in maintaining redox homeostasis and regulating metabolism in physio-pathological contexts, including cancer and inflammation (*50–52*). Asn restriction conferred robust effector function to CD8 T cells through enhancing the expression of inflammatory genes and rewiring metabolic programs to support nucleotide biosynthesis. Having shown that NRF2 is responsible for upregulating inflammatory genes under Asn restriction, we sought to determine if NRF2 is required for regulating the central carbon metabolism in CD8 T_eff_ cells following Asn restriction. While Asn restriction significantly reduced glucose/glutamine consumption and lactate/glutamate production in WT cells, NRF2 deletion partially reversed these changes (**Fig. 7A**). Similarly, NRF2 deletion abrogated the effects of Asn restriction on decreasing glycolysis and enhancing the PPP (**Fig. 7B**). Asn restriction could significantly increase basal and maximal respiration rate, mitochondrial mass, and DNA content in WT CD8 T cells. However, NRF2 deletion in CD8 T cells reversed most of the changes caused by Asn restriction (**Fig. 7C-D**), suggesting that NRF2 plays a broader role in controlling the central carbon metabolism upon Asn restriction.

**Figure 7.**
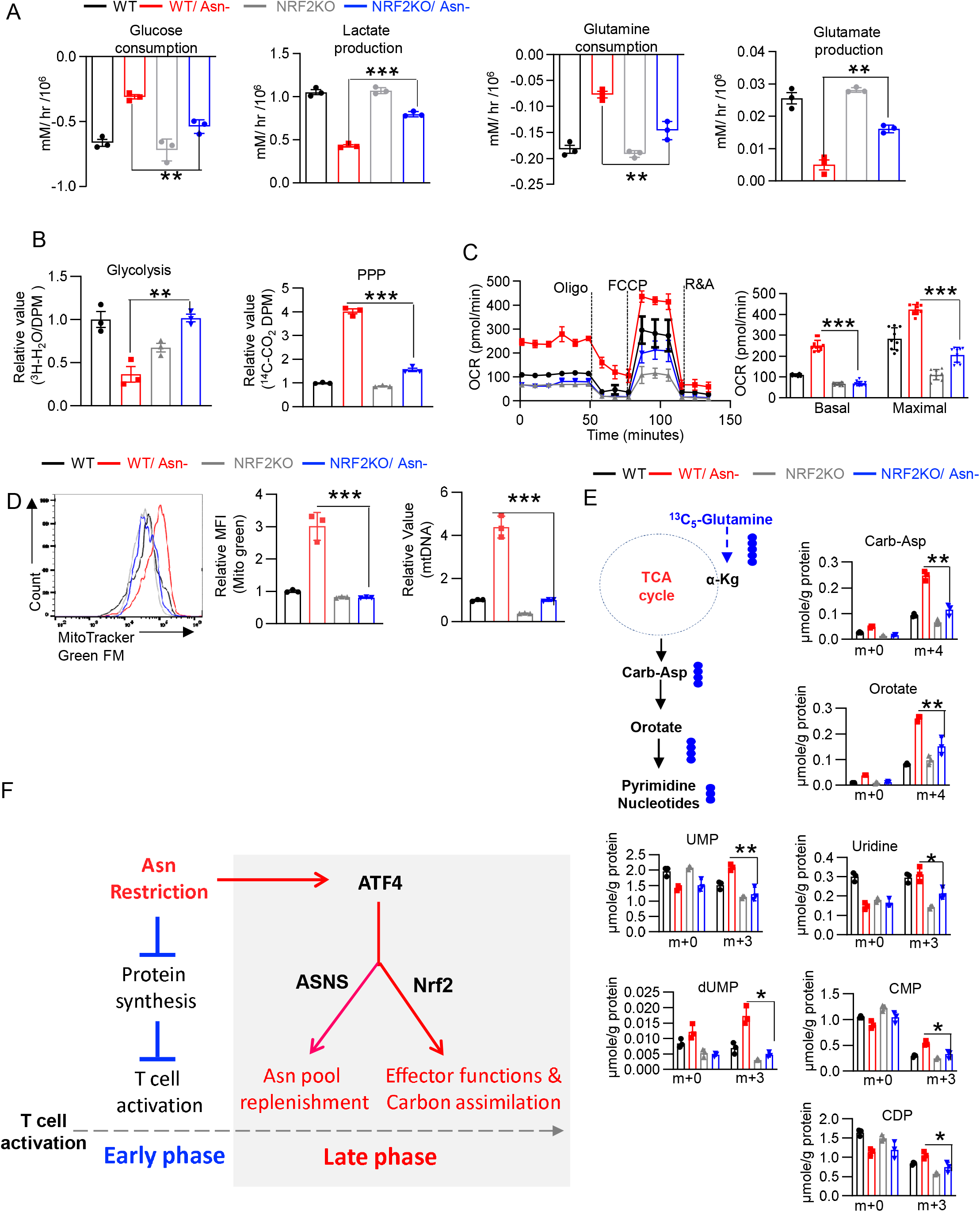
Asn restriction reprograms the central carbon metabolism through NRF2. A-D) CD8 T cells with the indicated genotype were activated in a medium with or without Asn. The indicated metabolites were quantified by the YSI bioanalyzer (A). The consumption and production were determined by calculating the difference between blank and spent medium (6hr incubation of T cells collected in the late phase/72hr) (n=3). The glycolysis activity and PPP activity (in the late phase/72hr) were determined by measuring ^3^H_2_O generated from D-[5-^3^H(N)]-glucose and ^14^CO_2_ generated from [1-^14^C] D-glucose, respectively (B) (n=3). OCR (in the late phase/72hr) was determined by Seahorse (C) (n=3). Mitochondria mass was determined by Mito Tracker green FM staining by flow cytometry, and mitochondrial DNA was quantified by qPCR (D) (n=3). Data from A-D are representative of 3 independent experiments. E) Diagram of [^13^C_5_]-Glutamine catabolism through entering the downstream TCA cycle and pyrimidine biosynthesis. ●: denoted the ^13^C label of all carbons of indicated metabolites derived from [^13^C_5_]-glutamine catabolism (left panel, E). Metabolites in cells (in the late phase/72hr) were analyzed by IC-UHR-FTMS (right panel, E), numbers on the X-axis represent those of ^13^C atoms in given metabolites, and numbers on the Y-axis represent the levels of the metabolites (μmole/g protein). Carb-Asp: N-carbamoyl-L-aspartate; UMP/UDP: uridine mono/diphosphate; dUMP deoxyuridine monophosphate; CMP/CDP: cytidine mono/diphosphate (n=3). Data are representative of one experiment. Error bars represent mean ± SD. *p < 0.05, **p < 0.01, *** P < 0.001, Student’s t-test. F) The conceptual model of ATF4/NRF2-dependent stress signaling response, optimized carbon assimilation, and robust effector function on CD8 T cell in the late phase.

While Asn restriction reduced the overall carbon consumption from glucose and glutamine (**Fig. 6B-C**), it significantly enhanced the de novo synthesis of nucleotides to promote cell proliferation (**Fig. 6H-I**). To determine if NRF2 is required for enhancing de novo synthesis of nucleotides upon Asn restriction. We supplied ^13^C_5_-glutamine as a metabolic tracer in T cell culture media and followed ^13^C incorporation into downstream metabolites. NRF2 deletion abolished the effect of Asn restriction in increasing fractional enrichment of ^13^C isotopologues in most metabolites (**Fig. 7E**). In line with previous studies that NRF2 regulated the metabolic genes of the PPP and nucleotide biosynthesis in cancer cells (*53, 54*), NRF2 deletion eliminated the effect of Asn restriction on inducing many of these metabolic genes (**Fig. S11A**). Collectively, our results indicated that Asn restriction engaged NRF2 to rewire the central carbon metabolism and promote de novo biosynthesis of nucleotide in CD8 T_eff_ cells.

## Discussion

Our studies showed that Asn restriction induced a biphasic response, consisting of the sequential action at two conceptually separated phases after CD8 T cell activation. Asn restriction suppressed CD8 T cell activation in the early phase by depriving intracellular Asn while rapidly engaging an ATF4/NRF2-dependent stress signaling response, conferring optimized carbon assimilation and robust effector function on CD8 T cell in the late phase (**Fig. 7F**). L-asparaginase is one of the first successful metabolic therapies and has been used in the clinic to treat acute lymphoblastic leukemia and lymphomas (*55, 56*). Asn-deficient media formulation can be incorporated into the current process of manufacturing CAR T cells (*57*). Our studies further implicate that Asn restriction is a promising and clinically relevant strategy to enhance cancer immunotherapy. Yet our findings on the Asn dependence of CD8 T_eff_ cells differed from two recent studies, which were also contradictory to each other (*58, 59*). While one study indicated that Asn restriction significantly inhibited CD8 T cell proliferation and effector functions, the other study showed that Asn restriction failed to compromise the CD8 T cell proliferation (*58, 59*). These discrepancies might be the result of some unknown technical issues.

The dependence of some but not other exogenous amino acids also reflects T cell’s capacity of preferentially engaging stress signaling to cope with specific environmental challenges. It has been recently shown that ATF4 deficiency rendered CD4 T cells sensitive to oxidative stress and incapable of eliciting proper immune responses (*60*). ATF4 mediates an integrated stress response in mammalian cells to various environmental insults, including nutrient restriction and oxidative stress (*47*). Similarly, NRF2 is a critical redox sensor that transcriptionally controls mammalian cells’ oxidative stress response (*61, 62*). Asn restriction, CD8 T cells developed a sophisticated acclimation program in response to Asn restriction. Cells switched to Asn de novo biosynthesis to replenish the intracellular Asn pool, reducing overall carbon consumption, enhancing nucleotide biosynthesis, and simultaneously eliciting robust effector functions. Such metabolic and phenotypical optimization was primarily achieved through engaging ATF4- and NRF2-dependent signaling in the early phase after T cell activation. ATF4 activation increased the expression of ASNS and NRF2, enabling Asn de novo biosynthesis and robust effector functions on CD8 T cells. While NRF2 was not required for promoting ASNS expression, NRF2 activation was necessary and sufficient to enhance the metabolic and functional fitness of CD8 T_eff_ cells.

The phenomena of incompletely oxidizing carbon sources and excreting intermediate metabolites is a shared metabolic feature in highly proliferating prokaryotic and eukaryotic cells (often referred to as overflow metabolism) (*63–65*). Our results revealed that CD8 T_eff_ cells excreted a wide range of metabolites, including nucleotide and their precursors, under the standard culture condition. It’s known that proliferating T cells only oxidize a small fraction of glucose to generate CO_2_ and energy while excreting most glucose-derived carbon into the extracellular compartment as lactate. Similarly, glutamine is only partially oxidized to CO_2,_ and a fraction of glutamine carbon is funneled into the extracellular compartment as glutamate and other metabolites in proliferating T cells (*66, 67*). Increased glycolysis and glutamine catabolism is a metabolic characteristic of activated T cells and is believed to be necessary for supporting their growth, proliferation, and effector function (*68–70*). However, we found that Asn restriction conferred robust proliferation and effector function to CD8 T cells, accompanied by an overall reduction of glucose and glutamine consumption. By contrast, Asn restriction markedly increased oxygen consumption and mitochondrial mass in CD8 T cells.

Mitochondria integrate the central carbon metabolism and critical cell signaling pathways to control fate-determining processes in mammalian cells. It has been recently reported that CD8 T_eff_ cells oxidized more glucose carbon in mitochondria in vivo than in vitro conditions (*71*). Similarly, reducing glutamine catabolism could promote oxidative phosphorylation and antitumoral activities of CD8 T_eff_ cells (*9*). In line with these findings, mitochondrial fitness is necessary for driving T cell proliferation, effector functions, and long-term persistence in vivo (*72–75*). However, the tumor microenvironment promotes T cell exhaustion and suppresses T cell-mediated antitumoral response through compromising mitochondrial biogenesis and function (*76–80*). CD8 T_eff_ cell metabolic fitness could be enhanced by optimizing carbon assimilation via increasing mitochondrial quality and quantity even when the overall carbon consumption was significantly reduced.

## Acknowledgments

We thank Dr. Xiaotong Song, Dr. Nicholas Restifo, and Dr. Williams Terence for reagents and advice. This work was supported by 1U01CA232488-01 from the National Institute of Health (Cancer Moonshot program), 2R01AI114581-06, and R01CA247941 from the National Institute of Health, V2014-001 from the V-Foundation, and 128436-RSG-15-180-01-LIB from the American Cancer Society (to RW). The Sanford Burnham Prebys Cancer Metabolism Core was supported by the SBP NCI Cancer Center Support Grant P30 CA030199. The Center for Environmental and Systems Biochemistry Core was supported in part by the Markey Cancer Center support grant P30CA177558

## Data availability statement

The RNA-seq datasets generated for this study can be found in the GEO accession GSE201870.

## Competing interests

All other authors declare no conflict of interest.

## Materials and Methods

### Mice

C57BL/6NJ, B6.129S4-Ifnγtm3.1Lky/J, Pmel^+^(B6.Cg-Thy1^a^/CyTg(TcraTcrb)8Rest/J), NRF2 KO (B6.129X1-Nfe2l2tm1Ywk/J), *Rag1*^-/-^ (B6.129S7-*Rag1^tm1Mom^*/J) and CD45.1 (B6.SJL-Ptprca Pepcb/BoyJ) were purchased from the Jackson Laboratory. NOD.CB17-Prkdcscid IL2rgtm1/BcgenHsd mice were purchased from Envigo. C57BL/6N-Asns tm1a(EUCOMM)Wtsi/H) mice and B6;129X1-Gt(ROSA)26Sortm2(ATF4)Myz/J were crossed with the CD4-Cre strain to generate T cell-specific ASNS KO and ATF4 overexpression (ATF-KI). ATF-KI mice were crossed with Pmel mice to generate Pmel^+^ ATF4 KI. ATF4^fl/fl^ mice were crossed with CD4-Cre strain to generate T cell-specific ATF4 KO (*81*). Gender and age-matched mice (6-12 weeks old) were used for the experiments. All mice were bred and kept in specific pathogen-free conditions at the Animal Center of Abigail Wexner Research Institute at Nationwide Children’s Hospital. Animal protocols were approved by the Institutional Animal Care and Use Committee of the Abigail Wexner Research Institute at Nationwide Children’s Hospital (IACUC; protocol number AR13-00055).

### T cells isolation and culture

Healthy donor buffy coats were obtained from the Central Ohio Region American Red Cross. The Institutional biosafety committee of the Abigail Wexner Research Institute at Nationwide Children’s Hospital approved this research. Mononuclear cells from peripheral blood (PBMCs) were isolated from buffy coats by Ficoll/Hypaque density gradient centrifugation. Isolated human mononuclear cells were stimulated with plate-bound antibodies in a medium containing human IL-2 (hIL-2). Plates were pre-coated with anti-human CD3 (hCD3) and anti-human CD28 (hCD28) overnight at 4°C. To generate GD2-CAR T cells, human PBMCs were activated using plate-bound anti-hCD3 and anti-hCD28 for 2 days, followed by GD2-CAR retroviral transduction and culture with hIL-2 for 7 days (*82*) Mouse T cells were enriched from spleen and lymph nodes by negative selection using a mouse CD8 naïve T cell isolation kit. For mouse T cell activation and proliferation assay, freshly isolated naïve CD8 T cells were activated by plate-bound antibodies and cultured with mouse IL-2 (mIL-2). The culture plates were pre-coated with anti-mouse CD3 (mCD3) and anti-mouse CD28 (mCD28) antibodies overnight at 4°C. For Pmel T cell activation and culture, splenocytes from Pmel transgenic mice were stimulated by human gp100 (hgp100) peptide and cultured with mIL-2 for 5 days, followed by a restimulation with mIL-2, soluble anti-mCD3, and anti-mCD28 for 2 days (*83*). To analyze intercellular cytokines and effector molecules, T cells were stimulated for 4hr with anti-mCD3, anti-mCD28, brefeldin A and monensin. In some experiments, Pmel T cells (2 days after activation) were transduced with NRF2-overexpressing retrovirus and cultured for 3 days (*83*). T cells were cultured with RPMI-1640 medium supplemented with 10% FBS, 2 mM L-glutamine, 100 units/mL penicillin, 100 μg/mL streptomycin, and 0.05 mM 2-mercaptoethanol at 37°C/5% CO_2_. For Asn restriction experiments, T cells were cultured with Asn-free RPMI 1640 medium supplemented with 10% (v/v) dialyzed fetal bovine serum (DFBS), 2 mM L-glutamine, 100 units/mL penicillin, 100 μg/mL streptomycin, and 0.05 mM 2-mercaptoethanol at 37°C/5% CO_2_. For culturing ATF4 KO T cells, Asn-free RPMI 1640 medium was supplemented with 10% (v/v) FBS, 2 mM L-glutamine, 100 units/mL penicillin, 100 μg/mL streptomycin, and 0.15 mM 2-mercaptoethanol as previously reported (*60*). The single amino acid-deficient medium was prepared by reconstituting RPMI 1640 amino acid-free basal medium with all but one (the corresponding amino acid). DFBS was made by dialyzing FBS against 100 volumes of distilled water (changed 5 times in 3 days) using dialysis cassettes (2K MWCO) at 4°C. Additional information on the concentration of antibodies, cytokines, and resources is listed in Table 1.

### Tumor cell line culture

Tumor cell lines were cultured with regular RPMI 1640 or Asn free RPMI 1640 medium supplemented with 10% (v/v) DFBS, 2 mM L-glutamine, and 100 units/mL penicillin, 100 μg/mL streptomycin at 37°C/5% CO_2_. CMT Keap1 KO cell line was generated by CRISPR by isolating a monoclonal cell population by limiting dilution.

### Adoptive cell transfer (ACT) for tumor immunotherapy

For the B16 melanoma ACT model, female C57BL/6 mice were implanted subcutaneously with 1x10^5^ cells of B16-gp100 in the flank, followed by sub-lethal irradiation (600 cGy) and intravenous injection of 1x10^6^ pmel T cells on day 7. For the LAN-1 xenograft, male Scid mice were implanted subcutaneously with 1.5 x 10^6^ LAN-1 tumor cells in the flank, followed by intravenous injection of 7 x 10^6^ GD2-CART cells on day 7. For the B16-F10 melanoma model, female C57BL/6 mice were implanted subcutaneously with 1x10^5^ cells of B16-gp100 in the flank, followed by intraperitoneal injection of anti-PDL1 antibody (twice per week). For the CMT Keap1 KO lung cancer model, female C57BL/6 mice were implanted subcutaneously with 1 x 10^5^ cells in the flank, followed by intraperitoneal injection of anti-PD1 and PDL1 antibodies (three-time per week). In some experiments, L-Asp was intraperitoneally administered every day. In all experiments, tumor sizes were measured daily until tumor size reached 2000 mm^3^. Tumor volume was calculated by length×width2×π/6.

### Adoptive cell transfer for homeostatic proliferation

For homeostatic proliferation in lymphopenic *Rag^-/-^* mice, naïve CD8 T cells isolated from donor mice were mixed with WT and KO cells at a 1:1 ratio and labeled with CFSE. Approximately 1×10^7^ cells in 100 μL PBS were intravenously injected into 6–8-week-old gender-matched host mice and treated with L-Asp for 3 days. Mice were sacrificed after 4 days, and lymph nodes and spleen were collected and processed to assess cell ratio and proliferation by flow cytometry analysis. Serum was collected from control and L-Asp treated mice, and metabolites in serum were quantified by GC-MS.

### Flow cytometry

For analyzing surface markers, cells were stained in PBS containing 2% (w/v) BSA and the indicated antibodies. In some experiments, T cells were stimulated by the cell stimulation cocktail (4hr at 37°C), followed by staining with antibodies for cell-surface protein. Then, cells were fixed and permeabilized by FoxP3 Fixation/Permeabilization solution, followed by staining with antibodies for intracellular proteins. Cell proliferation was determined by CFSE staining, and cell viability was evaluated by 7AAD staining. For analyzing DNA/RNA content, cells were stained with surface proteins, followed by fixation with 4% paraformaldehyde (30 mins at 4°C) and permeabilization with FoxP3 permeabilization solution. Then, cells were stained with 7AAD (5 min at room temperature) and pyronin-Y (30 min at room temperature) before being analyzed by a flow cytometer with PerCP and PE channels, respectively. For analyzing protein content, cells were incubated with O-propargyl-puromycin (OPP, 1 hr at 37°C), followed by fixation with FoxP3 permeabilization solution and 5 FAM-Azide stainings (0.5 hr at 37°C) according to protein synthesis assay kit. For analyzing the cell cycle profile, cells were incubated with BrdU (1 hr at 37°C), followed by cell surface staining, fixation, and permeabilization according to Phase-Flow Alexa Fluro 647 BrdU Kit. Flow cytometry data were acquired on Novocyte and were analyzed with FlowJo software. Additional information on flow cytometry antibodies, dilution, and resources is listed in Table 1.

### Real-time cytotoxicity assay

The target tumor cells (B16-gp100 mouse melanoma or LAN-1 human neuroblastoma cells) were used for determining the cytotoxic potential of Pmel T cell or GD2 CAR T using xCELLigence RTCA eSight. A 50-μl medium was added to E-Plates 96 for measurement of background values. Target cells were seeded in an additional 50 μl medium at a density of 3000 for B16-gp100 and 6000 for LAN-1 cells per well and cultured for 16-18 hr before adding T cells at an effector to target ratio (E: T) 5:1. Cell density and cell ratio were determined by previous titration experiments. Impedance-based measurements of the normalized cell index (CI) were performed every 15 min upon T cell addition. CI values were exported, and the percentage of lysis was calculated using RTCA Pro software.

### Quantification of cytokines in the cell culture supernatant

Pmel T cells were activated with hgp100 peptide for 4 days, followed by restimulation with anti-mCD3 and anti-mCD28 antibodies for 48hr. Cells were resuspended in fresh medium at a 2 x10^6^/mL density and cultured for 6 hr. Cytokine production in the cell culture supernatant was determined by ELISA. GD2 CART cells were resuspended in fresh medium at a 2 x10^6^/mL density and cultured for 6 hr. Cytokine production in the cell culture supernatant was determined by LEGENDplex™ human CD8/NK panel according to the manufacturer’s instructions. The data were collected by Novocyte and analyzed using LEGENDplex™ Data Analysis Software.

### Western blot analysis

Cells were harvested, lysed, and sonicated at 4°C in a lysis buffer (50 mM Tris-HCl, pH 7.4, 150 mM NaCl, 0.5% SDS, 5 mM sodium pyrophosphate, protease, and phosphatase inhibitor tablet). Cell lysates were centrifuged at 13,000 × g for 15 min to collect the supernatant. The protein concentrations were determined by the Pierce™ BCA Protein Assay kit. The samples were boiled in NuPAGE® LDS Sample Buffer and Reducing solution for 5 min. The proteins were separated by NuPAGE Protein Gels, transferred to PVDF membranes by the iBlot Gel Transfer Device, then incubated with primary antibodies followed by a corresponding secondary antibody. Immunoblots were developed with LI-COR. Detailed information for primary antibodies and secondary antibodies was provided in table 1.

### RNA extraction, qPCR, and RNAseq

Total RNA was isolated using the Quick-RNATM MiniPrep Kit and was reverse transcribed using random hexamer and M-MLV Reverse Transcriptase. BIO-RAD CFX284^TM^ Real-Time PCR Detection System was used for SYBR green-based quantitative PCR. The relative gene expression was determined by the comparative *CT* method, also referred to as the 2^−ΔΔ^*^CT^* method. The data were presented as the fold change in gene expression normalized to an internal reference gene (beta2-microglobulin) relative to the control (the first sample in the group). Fold change = 2^−ΔΔ^*^C^*T = [(*CT* gene of interest-*CT* internal reference)] sample A – [(*CT* gene of interest-*CT* internal _reference_)] sample B. Samples for each experimental condition were run in triplicated PCR reactions. Primer sequences were obtained from Primer Bank to detect target genes (Table 2).

For RNA sequencing analysis, total RNA was extracted using RNeasy Mini Kit and treated with DNase I according to the manufacturer’s instructions. After assessing the quality of total RNA using an Agilent 2100 Bioanalyzer and RNA Nanochip, 150 ng total RNA was treated to deplete ribosomal RNA (rRNA) using target-specific oligos combined with rRNA removal beads. Following rRNA removal, mRNA was fragmented and converted into double-stranded cDNA. Adaptor-ligated cDNA was amplified by limit cycle PCR. After library quality was determined via Agilent 4200 TapeStation and quantified by KAPA qPCR, approximately 60 million paired-end 150 bp sequence reads were generated on the Illumina HiSeq 4000 platform. Quality control and adapter trimming were accomplished using the FastQC (version 0.11.3) and Trim Galore (version 0.4.0) software packages. Trimmed reads were mapped to the Genome Reference Consortium GRCm38 (mm10) murine genome assembly using TopHat2 (version 2.1.0), and feature counts were generated using HTSeq (version 0.6.1). Statistical analysis for differential expression was performed using the DESeq2 package (version 1.16.1) in R, with the default Benjamini-Hochberg *p-value* adjustment method. The Ingenuity Pathway Analysis (IPA) software, the Gene Set Enrichment Analysis (GSEA) software, and the R Programming Language software were used to analyze gene signature and pathway enrichment.

### Quantification of metabolites in the cell culture supernatant

The naïve CD8 T cells were activated by plate-bound anti-mCD3 and anti-mCD28 antibodies for 72hr. Cells were resuspended in the indicated medium at the density of 2×10^6^/ml for 6hr. The levels of glucose, lactate, glutamine, and glutamate in blank medium (without cells) and spent medium were measured by the bioanalyzer (YSI 2900). The metabolite consumption and production were determined by calculating the difference between blank and spent medium. In some experiments, the arginine level was determined by L-Arginine Assay Kit.

### Radioactive tracer-based metabolic assays

The radioactive tracer-based metabolic assay was performed as described previously (*84*). Glycolysis was measured by the generation of ^3^H_2_O from [5-^3^H(N)] D-glucose, pentose phosphate pathway was measured by the generation of ^14^CO_2_ from [1-^14^C] D-glucose, glutamine oxidation activity was measured by the generation of ^14^CO_2_ from [U-^14^C]-glutamine. For assays generating ^14^CO_2_, 5×10^6^ T cells in 0.5 mL fresh medium were dispensed into 7mL glass vials with a PCR tube containing 50μL 0.2N NaOH glued on the sidewall. After adding 1 μCi radioactive tracer, the vials were capped using a screw cap with a rubber septum and incubated at 37 for 2 h. The assay was then stopped by injecting 100μL 5N HCL into the vial. Vials were kept at room temperate overnight to trap the ^14^CO_2_. The NaOH solution in the PCR tube was then transferred to scintillation vials containing 10 mL scintillation solution for counting. A cell-free sample containing the same amount of tracer was included as a background control. For assays generating ^3^H_2_O, 1μCi radioactive tracer was added to the suspension of one million cells in 0.5mL fresh medium in 48 wells, then incubated at 37 for 2 h. The assay was stopped by transferring samples to a 1.5 mL microcentrifuge tube containing 50μL 5N HCL, which was placed in a 20 mL scintillation vial containing 0.5 mL water. Then the vial was capped and sealed. ^3^H_2_O was separated from other radio-labeled metabolites overnight at room temperature by evaporation diffusion. The 1.5 mL microcentrifuge tube was removed, and a 10mL scintillation solution was added to the vial before counting. A cell-free sample containing 1μCi radioactive tracer was included as a background control. Radioactivity was measured by liquid scintillation counting.

### Stable isotope labeling

Experiments in Fig.S5C-D: For isotope labeling of T cells in the early phase after activation, 2 ×10^6^ cells /mL naïve CD8 T cells were activated by plate-bound anti-mCD3 and anti-mCD28 antibodies in a conditional medium containing 4 mM ^13^C_5_-Glutamine for 12hr. For isotope labeling of T cells in the late phase after activation, naïve CD8 T cells were activated by plate-bound anti-mCD3 and anti-mCD28 antibodies for 72hr. Then cells are reseeded at a density of 2 ×10^6^ cells /mL in a conditional medium containing 4 mM ^13^C_5_-Glutamine and cultured for 6hr. Samples were collected and washed 3 times with PBS before being snap-frozen.

Experiments in Fig. 3C and 6H: Naïve CD8 T cells were activated by plate-bound anti-mCD3 and anti-mCD28 antibodies for 36hr. Then, cells were reseeded at a density of 2 ×10^6^ cells /mL in a conditional medium containing 4 mM ^13^C_5_-Glutamine for 24hr. Samples were collected and washed 3 times with PBS before being snap-frozen.

Experiments in Fig. 7E: Naïve CD8 T cells were activated by plate-bound anti-mCD3 and anti-mCD28 antibodies for 72hr. Then, cells were reseeded at the density of 2 ×10^6^ cells /mL in a conditional medium containing 4 mM ^13^C_5_-Glutamine for 6hr. Samples were collected and washed 3 times with PBS before being snap-frozen.

### Gas chromatography-mass spectrometry (GC-MS)-based analysis

GC-MS was performed as previously described (*85*). Cell pellets were resuspended in 0.45 mL – 20°C methanol/ water (1:1 v/v) containing 20 µM L-norvaline as the internal standard. Further extraction was performed by adding 0.225 mL chloroform, vortexing, and centrifugation at 15,000 ×*g* for 5 min at 4°C. The upper aqueous phase was evaporated under vacuum using a Speedvac centrifugal evaporator. Separate tubes containing varying amounts of standards were evaporated. Dried samples and standards were dissolved in 30 μL isobutylhydroxylamine hydrochloride in pyridine and incubated for 20 min at 80°C. An equal volume of N-tertbutyldimethylsilyl-N-methyltrifluoroacetamide (MTBSTFA) was added and incubated for 60 min at 80°C. After derivatization, samples and standards were analyzed by GC-MS using a Rxi-5ms column (15 m × 0.25 i.d. × 0.25 μM, Restek) installed in a Shimadzu QP-2010 Plus gas chromatograph-mass spectrometer (GC-MS). The GC-MS was programmed with an injection temperature of 250°C, 1.0 µL injection volume, and a split ratio of 1/10. The GC oven temperature was initially 130°C for 4 min, rising to 250°C at 6°C/min and to 280°C at 60°C/min with a final hold at this temperature for 2 min. GC flow rate, with helium as the carrier gas, was 50 cm/s. The GC-MS interface temperature was 300°C, and (electron impact) ion source temperature was 200°C, with 70 eV ionization voltage. Fractional labeling from ^13^C substrates and mass isotopomer distributions were calculated as described (*85*). Data from standards were used to construct standard curves in MetaQuant (*86*), from which metabolite amounts in samples were calculated. Metabolite amounts were corrected for recovery of the internal standard and for ^13^C labeling to yield total (labeled and unlabeled) quantities in nmol per sample and then adjusted to protein.

### Ion chromatography-ultra high resolution-fourier transform mass spectrometry (IC-UHR-FTMS)-based analysis

The frozen cell pellets were homogenized in 60% cold CH3CN in a ball mill for denaturing proteins and optimizing extraction. Polar metabolites were extracted by the solvent partitioning method with a final CH3CN: H2O: CHCl3 (2:1.5:1, v/v) ratio, as described previously (*87*). The polar extracts were reconstituted in nanopure water before analysis on a Dionex ICS-5000+ ion chromatography system interfaced with a Thermo Fusion Orbitrap Tribrid mass spectrometer as previously described (*88*) using a m/z scan range of 80-700. Peak areas were integrated and exported to Excel via the Thermo TraceFinder (version 3.3) software package before natural abundance correction (*89*). The isotopologue distributions of metabolites were calculated as the mole fractions as previously described (*90*). The number of moles of each metabolite was determined by calibrating the natural abundance-corrected signal against authentic external standards. The amount was normalized to the amount of extracted protein and is reported in µmol/g protein.

### Liquid chromatography-mass spectrometry (LC-MS)-based analysis

Sample preparation and analysis were carried out as described previously at Metabolon, Inc (*91*). In brief, sample preparation involved protein precipitation and removal with methanol, shaking, and centrifugation. The resulting extracts were profiled on an accurate mass global metabolomics platform consisting of multiple arms differing by chromatography methods and mass spectrometry ionization modes to achieve broad coverage of compounds differing by physiochemical properties such as mass, charge, chromatographic separation, and ionization behavior. Metabolites were identified by automated comparison of the ion features in the experimental samples to a reference library of chemical standard entries that included retention time, molecular weight (*m/z*), preferred adducts, and in-source fragments as well as associated MS spectra and were curated by visual inspection for quality control using software developed at Metabolon. Metaboanalyst was used to range-scale data and provide pathway analysis of metabolites significantly changed (P<0.05).

### Oxygen consumption rate (OCR)

OCR was determined by seahorse XFe96 Analyzer according to the manufacturer’s instruction. Briefly, naïve CD8 T cells were activated by plate-bound anti-mCD3 and anti-mCD28 antibodies for 72hr. 1 ×10^5^ cells were harvested, washed by PBS, and suspended in 50 µL assay medium (Seahorse XF RPMI Assay Medium) containing 10 mM glucose, 2 mM glutamine, and 1 mM pyruvate, and seeded each well in a Poly-D-Lysine pre-coated XF96 Cell Culture Microplates. Then, the plate was centrifuged at 200 ×*g* for 2 min in a zero-braking setting to immobilize cells, followed by adding a 130 µL assay medium and incubating in a non-CO_2_ incubator for 30 mins. OCR was measured under basal conditions and in response to oligomycin, fluoro-carbonyl cyanide phenylhydrazone (FCCP), or 100 nM rotenone with 1 µM antimycin A. Data analysis was performed using the Seahorse Wave Software.

### ATP levels measurement

The ATP levels were measured according to the manufacturer’s instructions of the CellTiter-Glo^®^ 2.0 Assay kit. Briefly, 1×10^5^ cells were suspended with 50uL PBS in 96-well white-walled tissue culture plates, and an equal volume of CellTiter-Glo 2.0 reagent was added to each well. The plate was put on a shaker for gentle mixing and incubated at room temperature for 30 minutes. The luminescence was measured by the Microplate Reader.

### Statistical Analysis

Statistical analysis was conducted using the GraphPad Prism software. P values were calculated with Two-way ANOVA for in vivo experiments. Kaplan-Meier curves and the corresponding log-rank test was used to evaluate the statistical differences between groups in survival studies. Paired and unpaired two-tail Student’s t-tests were used to assess differences in other experiments. R Programming Language software was used for Metabolon and RNA-seq data analysis. P values less than 0.05 were considered significant, with p < 0.05, p < 0.01, p < 0.001 indicated as *, **, and ***, respectively, ns indicated no significant differences.

---

**Supplemental Figure 1.**
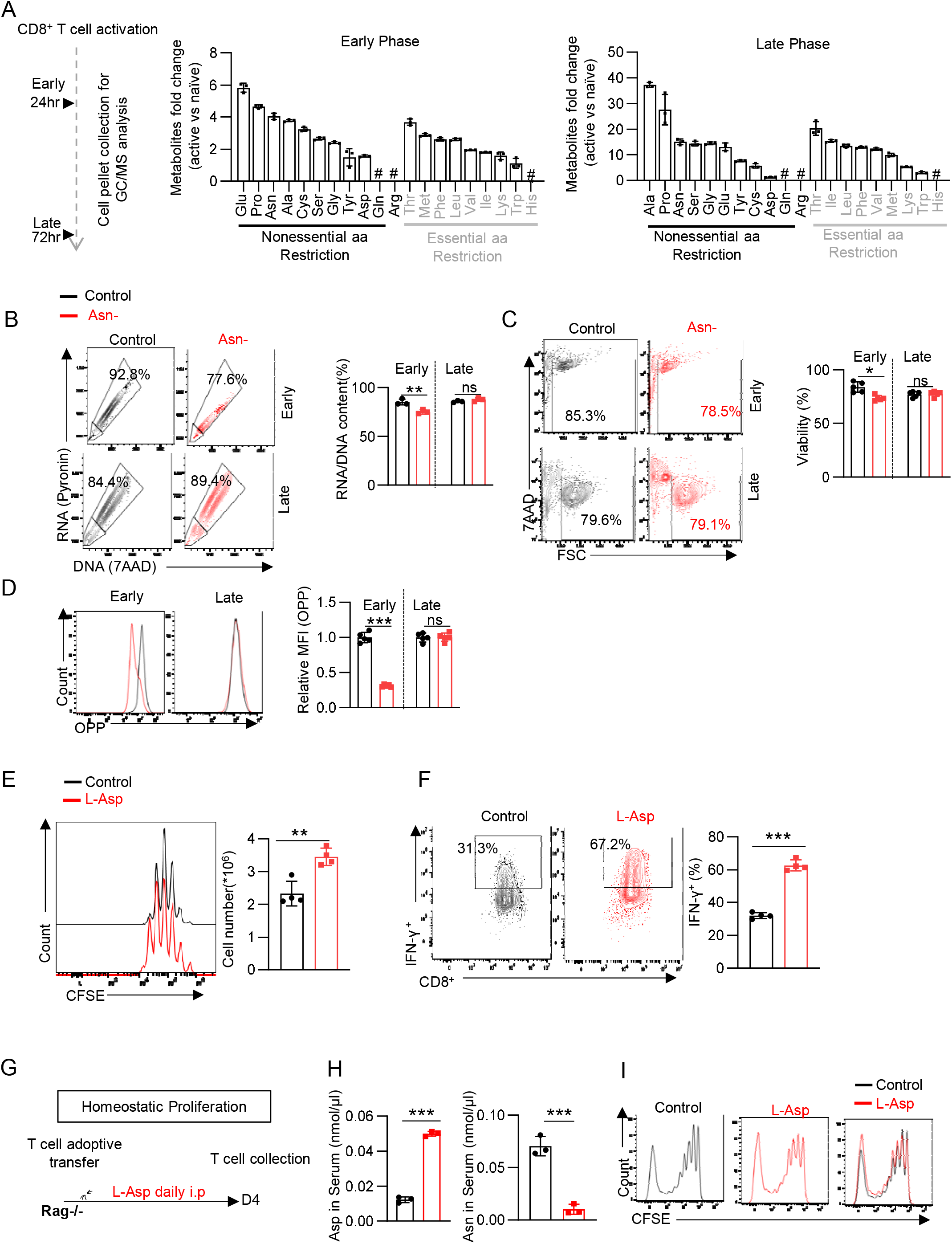
Asn restriction on CD8 T cells induces a biphasic response. A) Schematic diagram of measuring amino acid consumption and production of T cells in the early (24hr) and late phase (72hr) after activation (left). Indicated amino acids in the cell pellet collected from T cells in the early (middle) and the late phase (right) were quantified by GC-MS. Y-axis represents the fold change of active T cells over naïve T cells. # Indicates amino acids that were not quantified due to technical limitations (n=3). Data are representative of 2 independent experiments. B-D) CD8 T cells were activated in a medium with or without Asn and were collected in the early phase (24hr) and late phase (72hr). The DNA/RNA content (B) (n=3), cell viability (C) (n=5), and protein content (D) (n=5) was determined by flow cytometry. Data from B-D are representative of 3 independent experiments. E-F) CD8 T cells were activated in a medium with or without asparaginase (L-Asp) and were collected in the late phase (72hr). Cell proliferation was determined by CFSE staining by flow cytometry, and the cell number (E) was determined by a cell counter (n=4). The expression of indicated cytokine was determined by flow cytometry (F) (n=4). Data are representative of 3 independent experiments. G) Schematic diagram of in vivo proliferation. (H) Mouse serum Asp and Asn levels were quantified by GC-MS (n=3). Data are representative of 2 independent experiments. I) Cell proliferation was determined by CFSE staining by flow cytometry. Data are representative of 3 independent experiments. Error bars represent mean ± SD. *p < 0.05, **p < 0.01, *** P < 0.001, Student’s t-test.

**Supplemental Figure 2.**
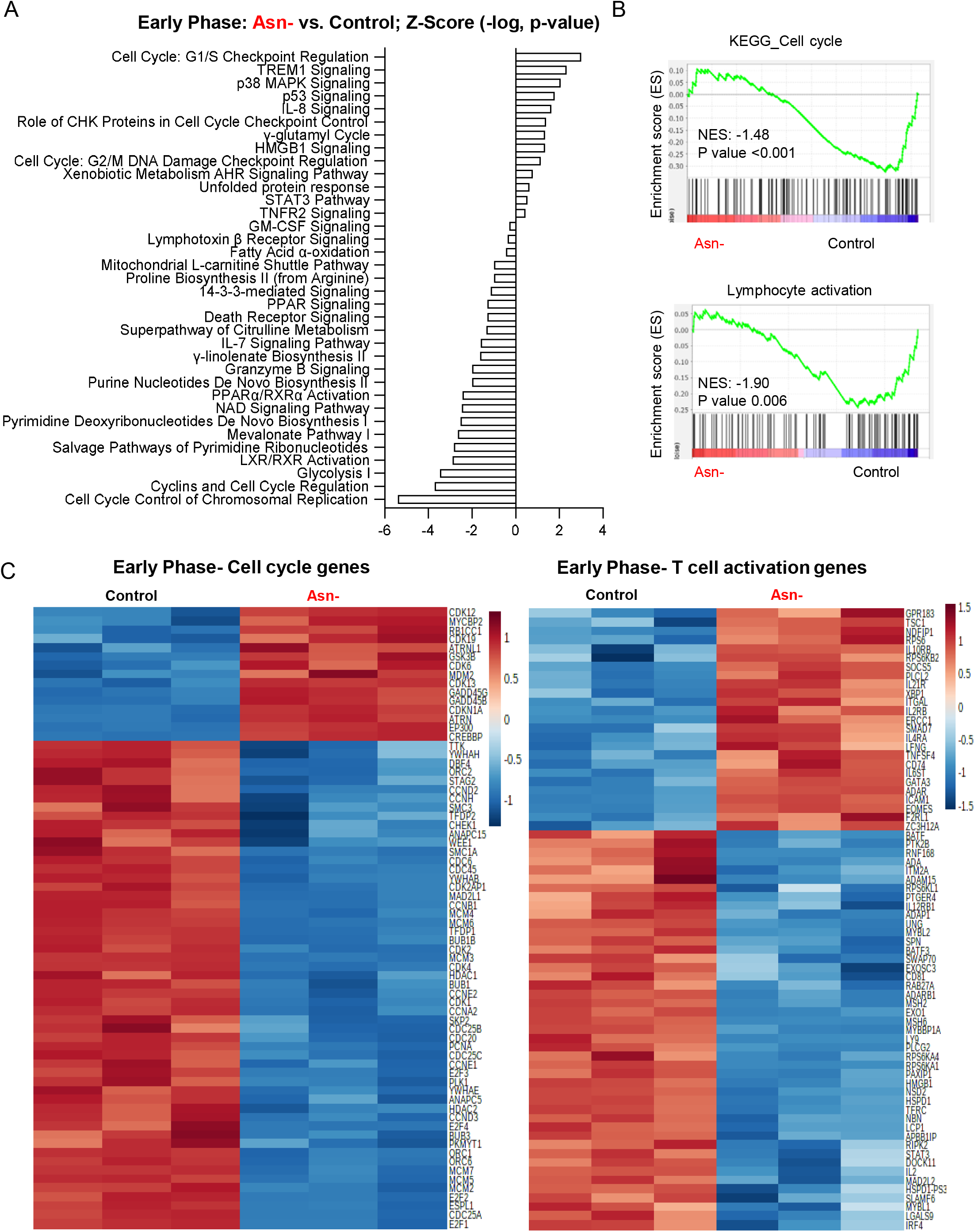
Asn restriction suppresses the expression of the cell cycle- and activation-related genes in the early phase after T cell activation. A-C) RNA seq was performed with CD8 T cells that have been activated in a medium with or without Asn for 24hr (the early phase). Differently expressed gene signatures were determined by the IPA analysis and listed according to their z score (A). Cell cycle and lymphocyte activation-associated gene sets were analyzed by GSEA (B). Heatmap depicting top 70 differentially expressed cell cycle genes (left) T cell activation genes (right) (C) (n=3). Data are representative of one experiment.

**Supplemental Figure 3.**
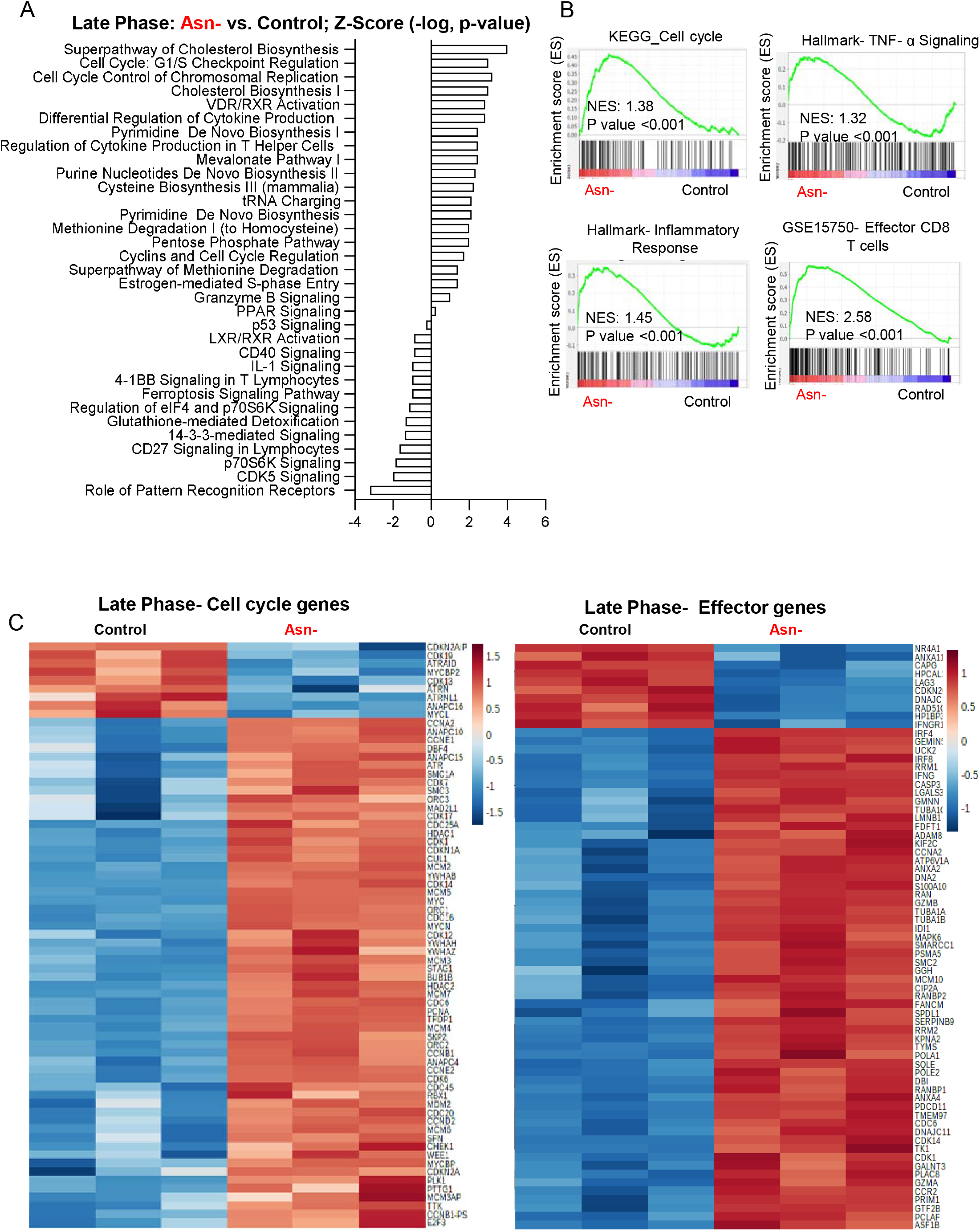
Asn restriction induces the expression of pro-inflammatory genes in the late phase after T cell activation. A-C) RNA seq was performed with CD8 T cells that have been activated in a medium with or without Asn for 72hr (the late phase). Differently expressed gene signatures were determined by the IPA analysis and listed according to their z score (A). Cell cycle, TNF-α, inflammatory response, and effector T cell function associated gene sets were analyzed by GSEA (B). Heatmap depicting top 70 differentially expressed cell cycle genes (left) and effector markers (right) (C) (n=3). Data are representative of one experiment.

**Supplemental Figure 4.**
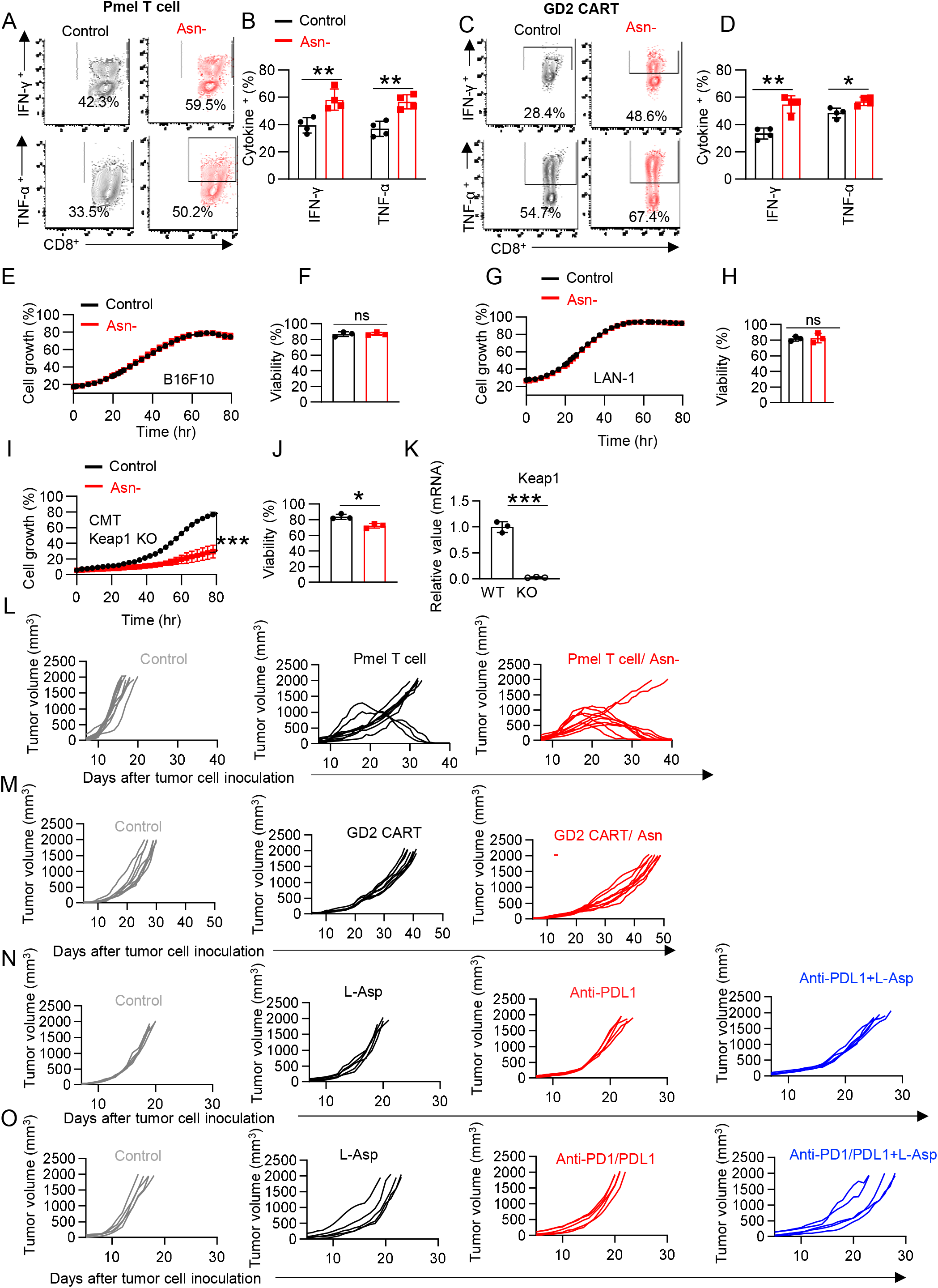
Asn restriction differentially affects T cells and tumor cells. A-D) The indicated cytokines of cells (described in Fig. 2A-B) were quantified by intracellular staining by flow cytometry (n=4). Data are representative of 3 independent experiments. E-J) B16F10, LAN-1, and CMT Keap1 KO cells were cultured in a medium with or without Asn. Cell growth (E, G, and I) was determined by live-cell imaging analysis (IncuCyte ZOOM™) (n=3). Cell viability (F, H, and J) was determined by 7AAD staining by flow cytometry (n=3). Data are representative of 3 independent experiments. K) Keap1 mRNA expression in CMT lung cancer cells was determined by qPCR (n=3). Data are representative of 2 independent experiments. Error bars represent mean ± SD. *p < 0.05, **p < 0.01, ns, not significant, Student’s t-test. L-O) Tumor growth of indicated groups (L represents Fig. 2F, M represents Fig. 2H, N represents Fig. 2J, and O represents Fig. 2L).

**Supplemental Figure 5.**
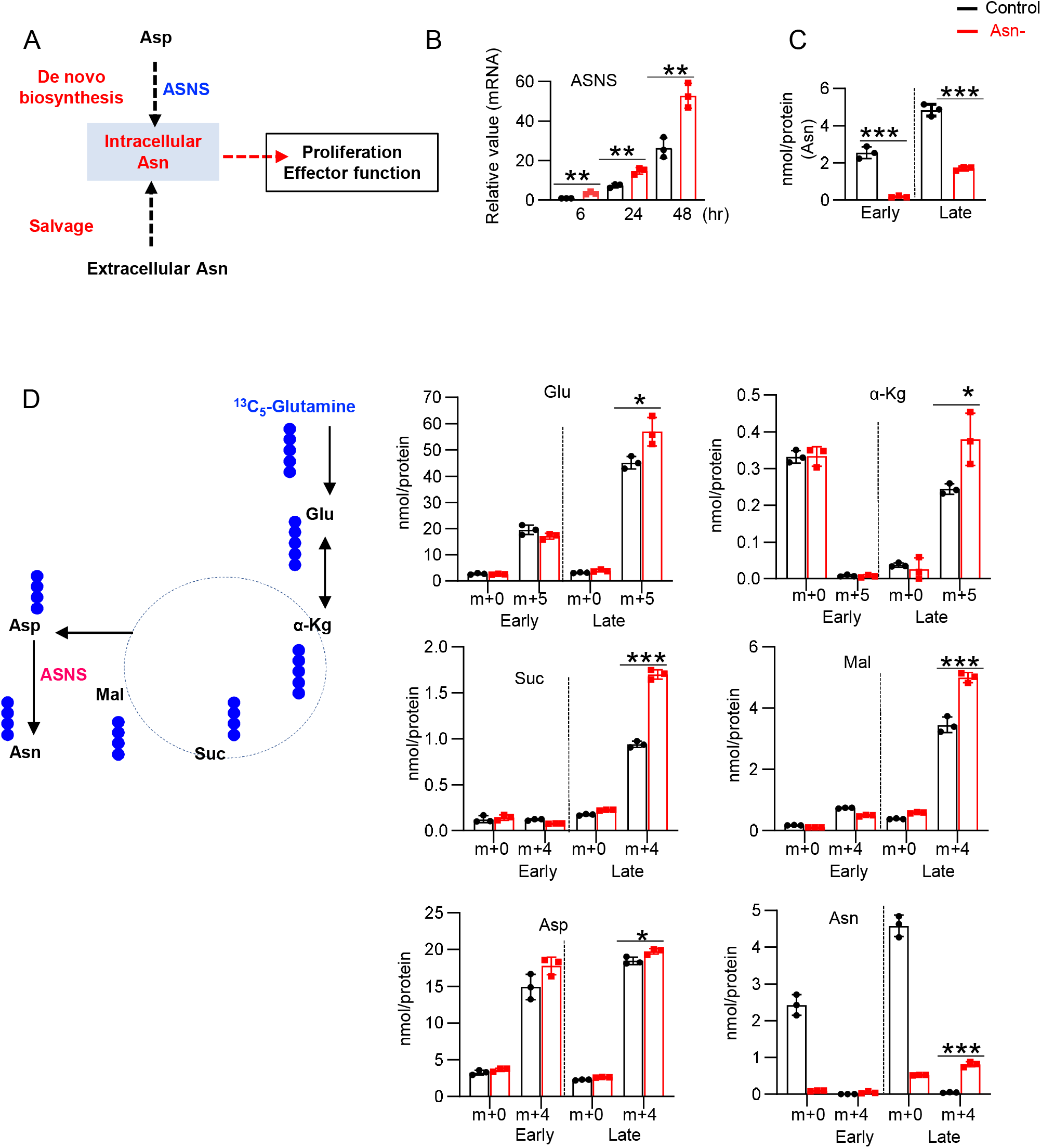
Asn restriction promotes Asn de novo biosynthesis in the late but not the early phase after T cell activation. A) Conceptual diagram of maintaining intracellular Asn pool in T cells. B) ASNS mRNA expression in the indicated groups was determined by qPCR (n=3). Data are representative of 2 independent experiments. C) The level of Asn in the indicated groups was determined by GC-MS (n=3). D) Diagram of [^13^C_5_]-Glutamine catabolism through entering the downstream TCA cycle and Asn biosynthesis. ●: denoted the ^13^C label of all carbons of indicated metabolites derived from [^13^C_5_]-glutamine catabolism (left panel). Metabolites in the indicated groups were analyzed by GC-MS (right panel), numbers on the X-axis represent those of ^13^C atoms in given metabolites, and numbers on the Y-axis represent the levels of the metabolites (nmol/ protein). Glu: glutamate, α-Kg: α-ketoglutarate, Suc: succinate, Mal: malate, Asp: aspartate, Asn: asparagine, (n=3). Data are representative of one experiment. Error bars represent mean ± SD. *p < 0.05, **p < 0.01, *** P < 0.001, Student’s t-test.

**Supplemental Figure 6.**
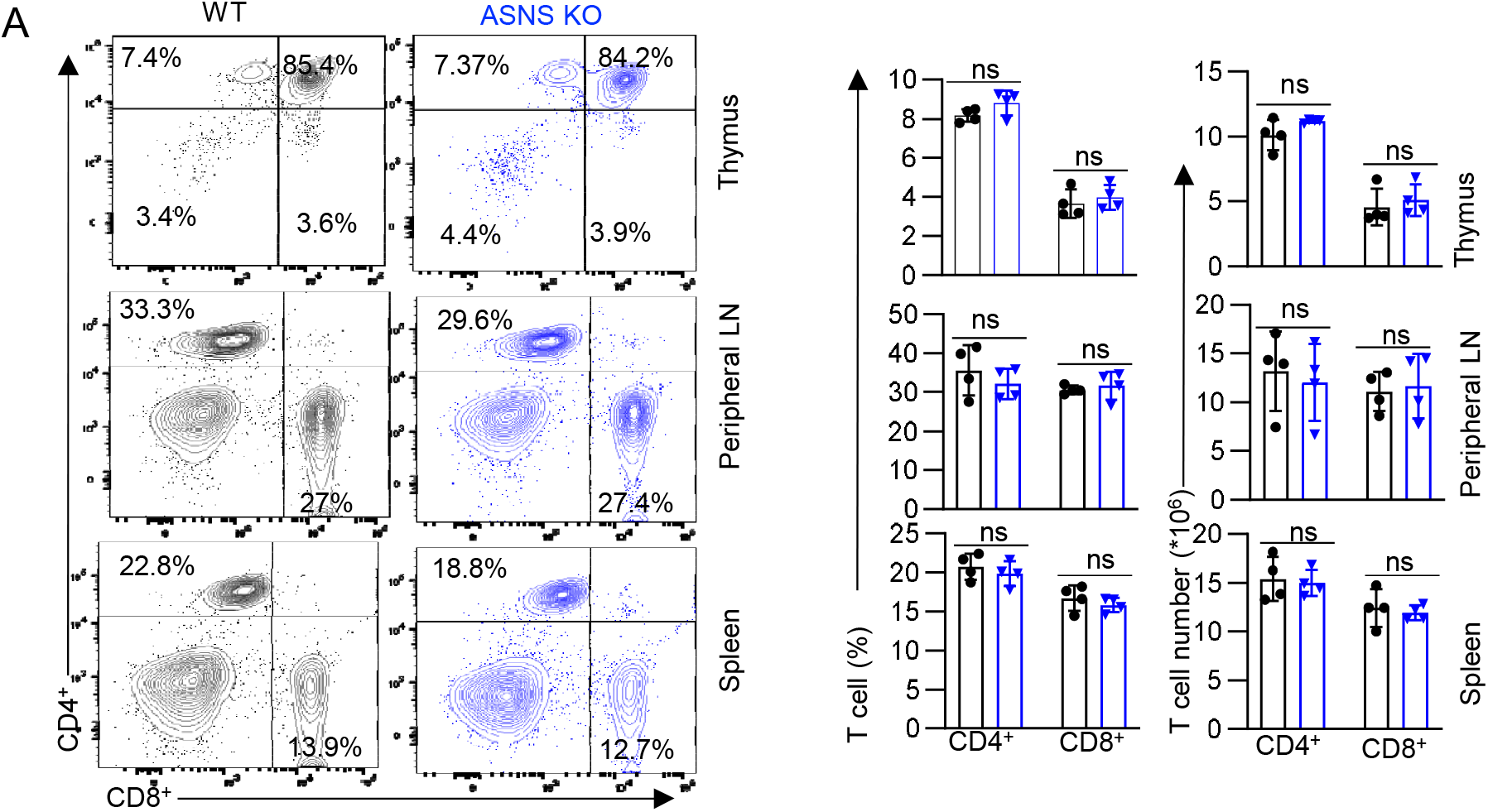
ASNS is dispensable for normal T cell development after the double-positive stage. A) Representative flow plots (left) and quantification of CD4 and CD8 distribution in the thymus, peripheral lymph nodes, and spleen (right) (n=4), Data are representative of 2 independent experiments. Error bars represent mean ± SD. ns, not significant, Student’s t-test.

**Supplemental Figure 7.**
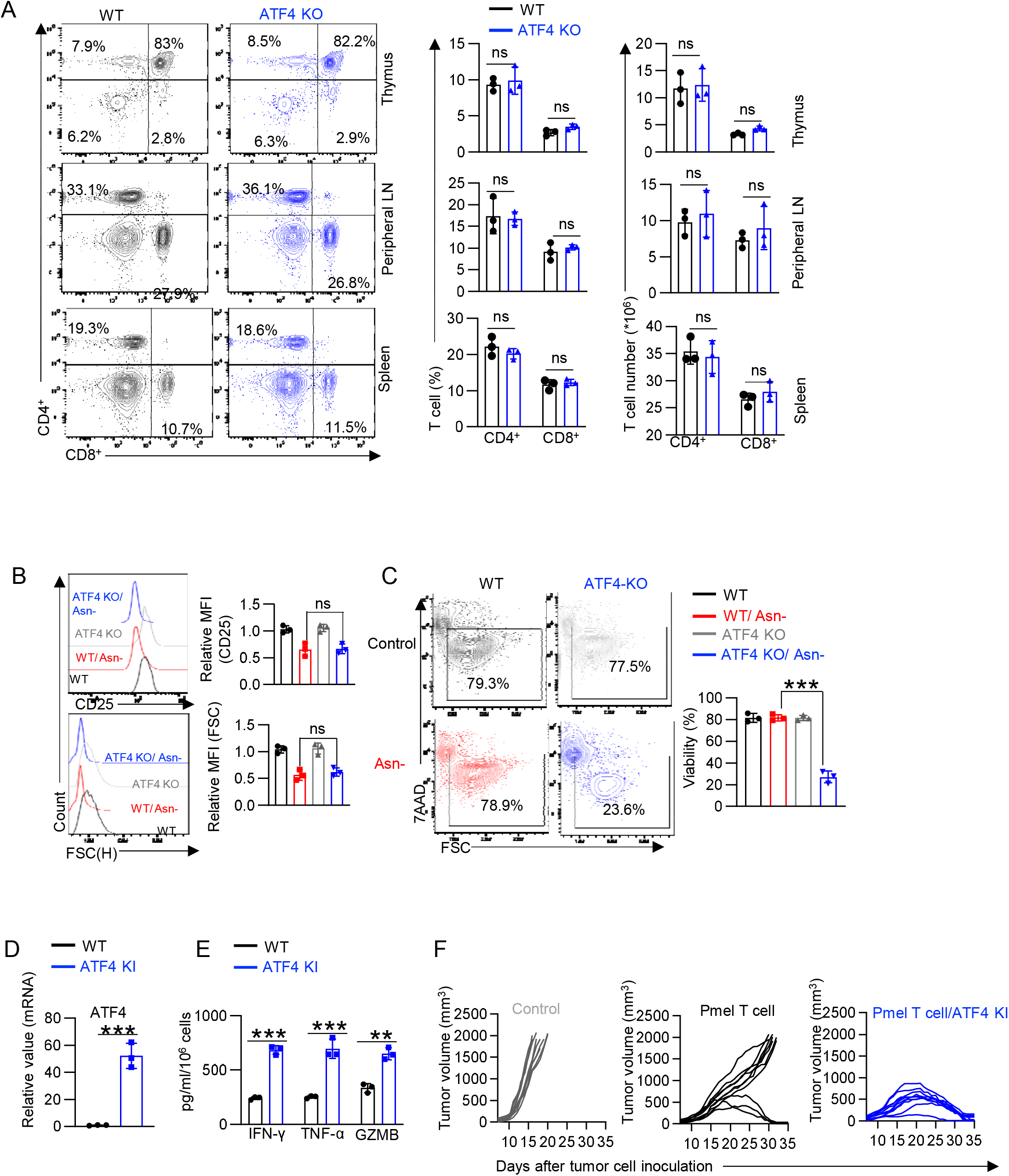
ATF4 regulates T cell development in vivo and T cell function in vitro. A) Representative flow plots (left) and quantification of CD4 and CD8 distribution (right) in the thymus, peripheral lymph nodes, and spleen (n=3). Data are representative of 2 independent experiments. B-C) CD8 T cells with the indicated genotype were activated in a medium with or without Asn and collected at 24hr (the early phase, B) and 72hr (the late phase, C). Cell surface expression of CD25, cell size (FSC) (B) (n=3), cell viability (C) (n=3) was determined by flow cytometry. Data are representative of 2 independent experiments. D-E) The level of the indicated mRNA was determined by qPCR (D) (n=3). The indicated protein in the cell culture medium collected 5 days after activation was quantified by ELISA (E) (n=3). Data are representative of 2 independent experiments. Error bars represent mean ± SD. **p < 0.01, *** P < 0.001, ns, not significant, Student’s t-test. F) Tumor growth of indicated groups of data presented in Fig. 4L.

**Supplemental Figure 8.**
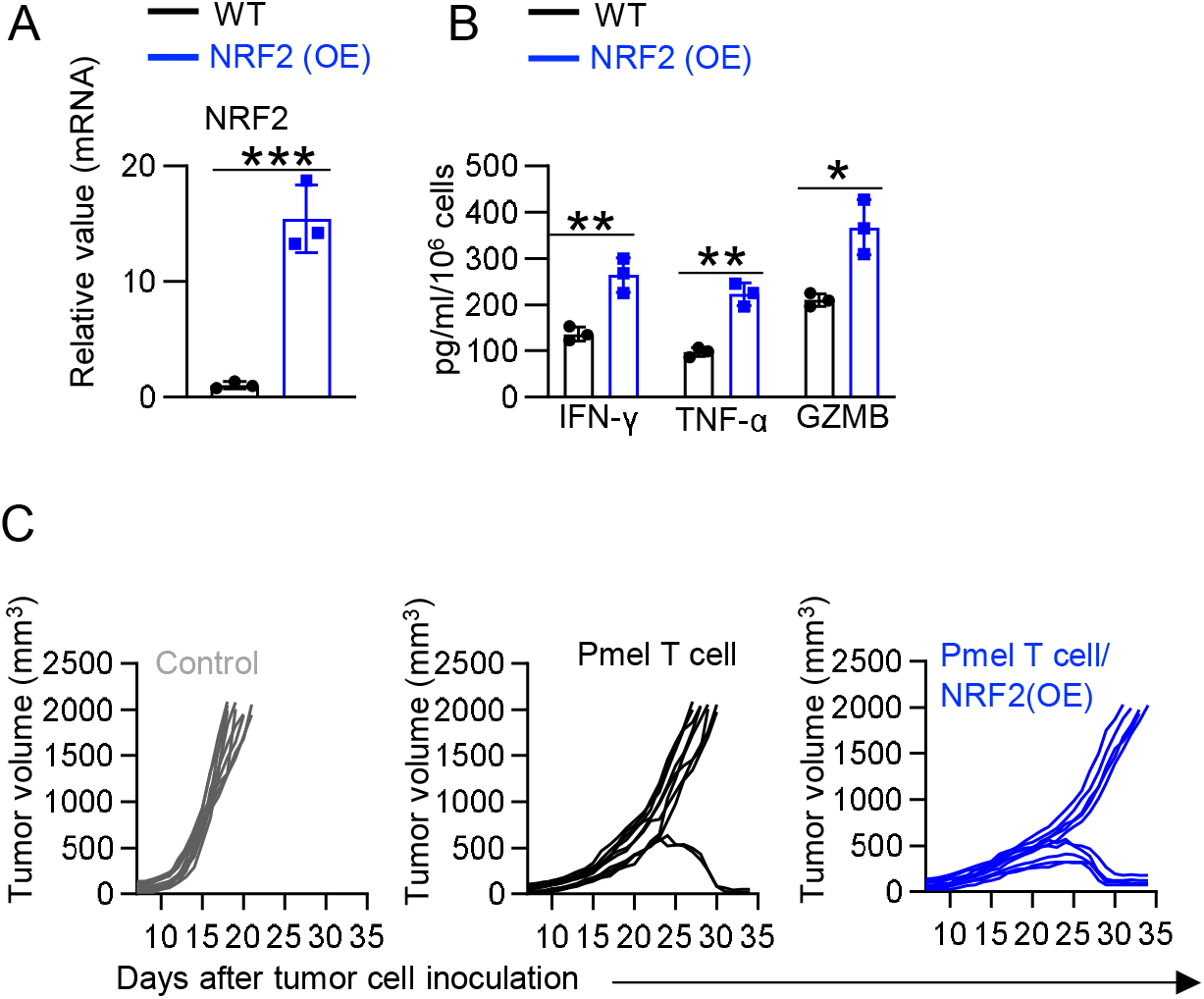
NRF2 control CD8 T cell effector function. A) The level of the indicated mRNA was determined by qPCR (n=3). B) The indicated protein in the cell culture medium collected 5 days after activation was quantified by ELISA (n=3). Data from A-B are representative of 2 independent experiments. Error bars represent mean ± SD. **p < 0.01, *** P < 0.001, Student’s t-test. C) Tumor growth of indicated groups of data presented in Fig. 5H.

**Supplemental Figure 9.**
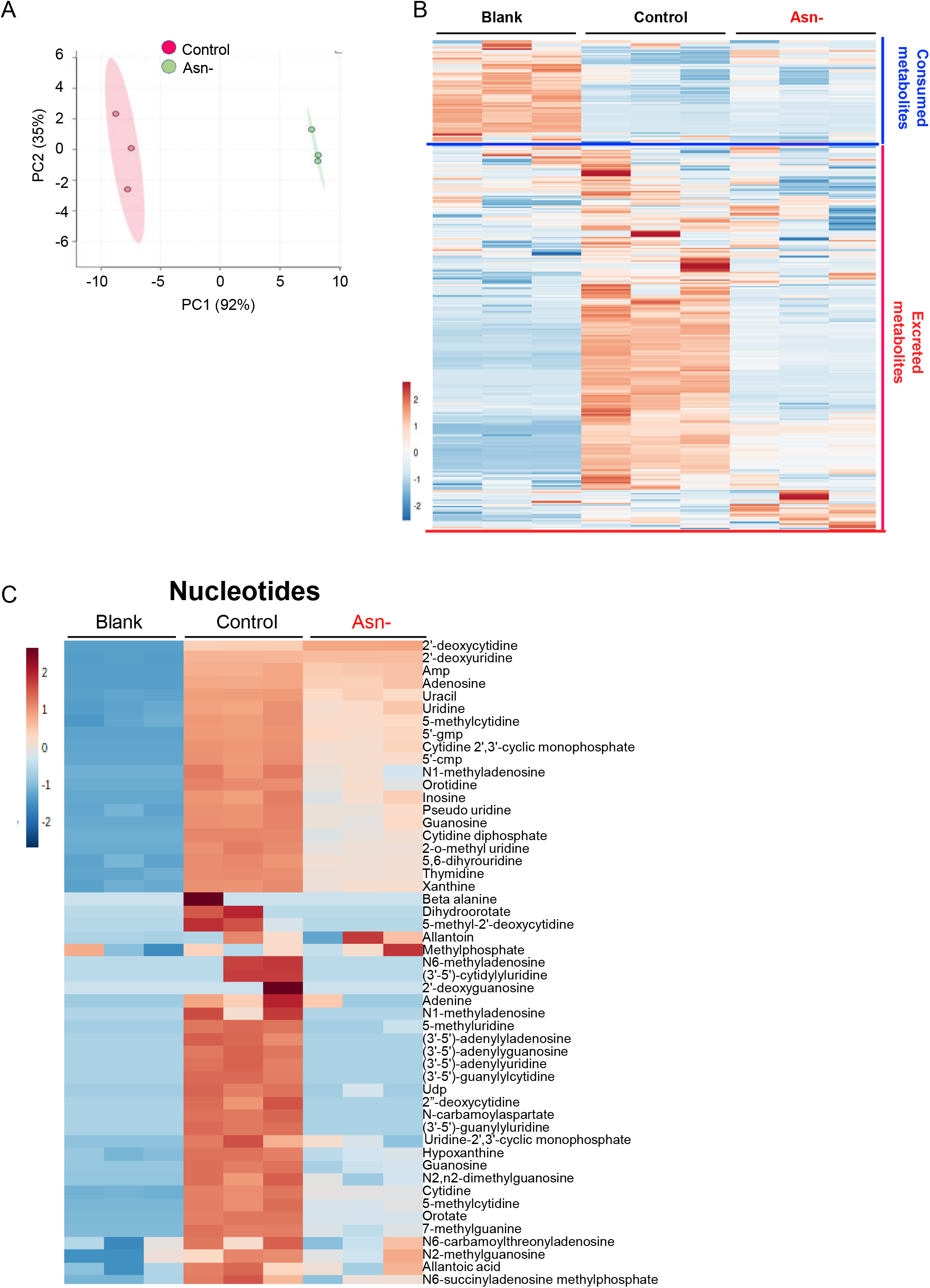
Asn restriction rewires the central carbon metabolism of CD8 T cells. A-C) The extracellular metabolome of indicated groups was determined by LC-MS (Fig. 6A). Principal component analysis (PCA) of the extracellular metabolome of indicated groups (A). Heatmap depicting differential production and consumption of extracellular metabolites (B). Heatmap depicting the level of indicated nucleotide metabolites in the indicated medium (C) (n=3). Data are representative of one experiment.

**Supplemental Figure 10.**
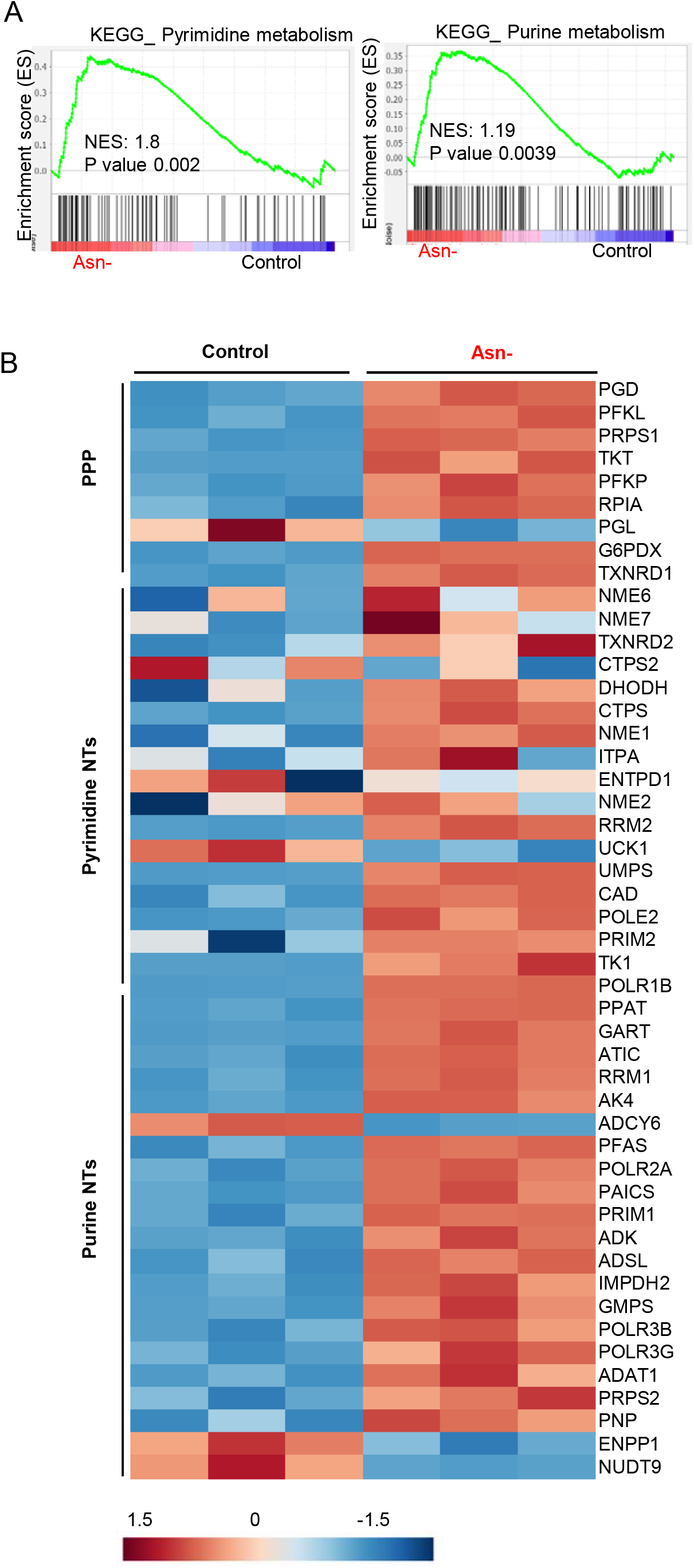
Asn restriction increases the expression of genes involved in nucleotide metabolism. A-B) RNA seq was performed with CD8 T cells that have been activated in a medium with or without Asn for 72hr (the late phase). Gene set enrichment analysis (GSEA) (A) and heatmap depicting differentially expressed metabolic genes (B) (n=3). Data are representative of one experiment.

**Supplemental Figure 11:**
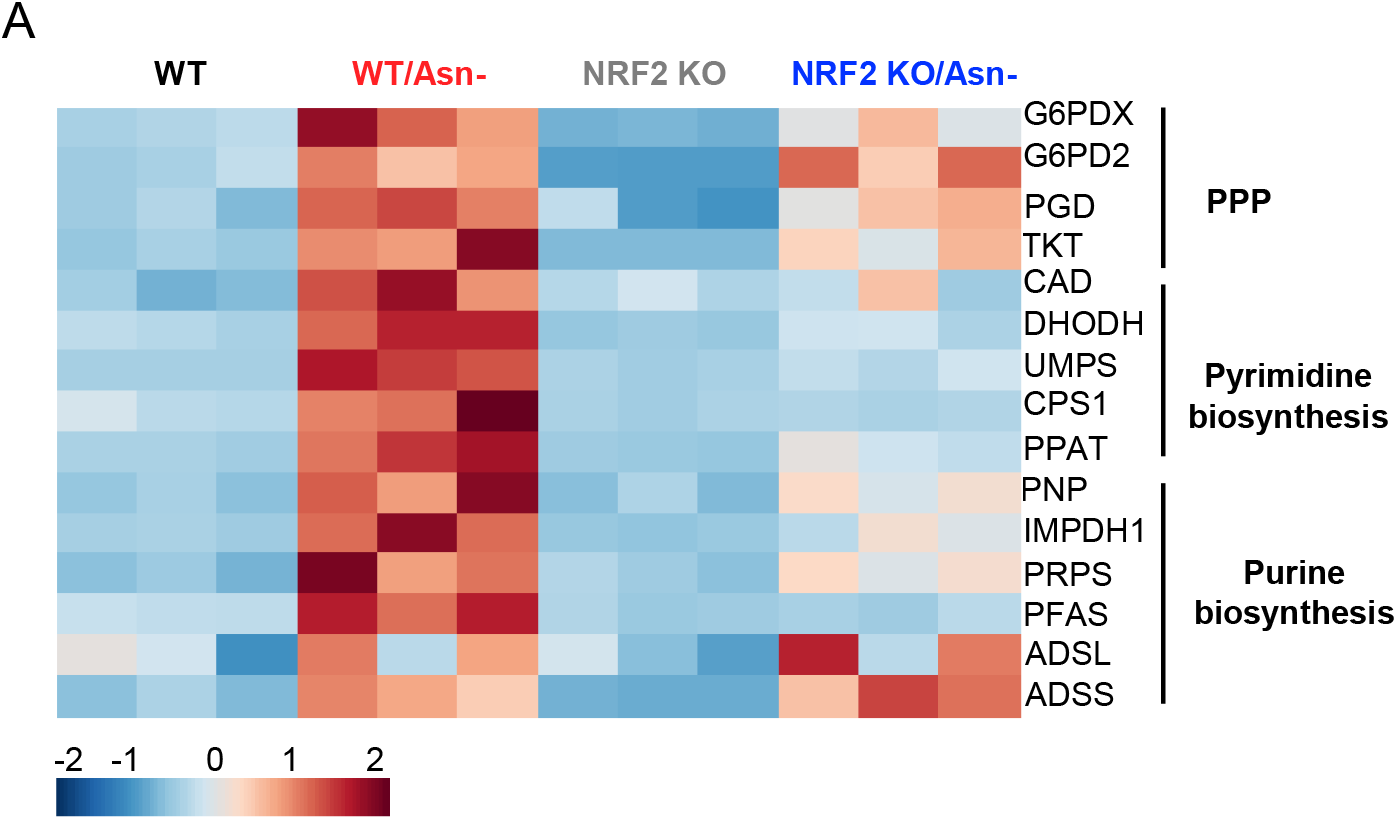
NRF2 is required for upregulating metabolic genes conferred by Asn restriction. A) The level of the indicated mRNA in cells collected 72hr after activation was determined by qPCR and depicted by heatmap (C) (n=3). Data are representative of 2 independent experiments.

## Notes

### Competing Interest Statement

The authors have declared no competing interest.

